# Comparative phylogeography of lizards (Squamata: Phrynosomatidae) in Baja California and expansion of *Callisaurus draconoides* within the North American deserts

**DOI:** 10.1101/2025.04.29.650089

**Authors:** Andrew D. Gottscho, Bradford D. Hollingsworth, Julio Lemos Espinal, Adam D. Leaché, Tod W. Reeder, Kevin de Queiroz

## Abstract

We examined the comparative phylogeography of five co-distributed lizard complexes (representing four genera within Phrynosomatidae) along the Baja California Peninsula (BCP) in the context of the region’s complex geological history. Double-digest restriction-associated-DNA (ddRAD) sequencing was used to collect genome-wide sequence data for 228 peninsular lizards. Using phylogenetic analyses of concatenated loci and population admixture analysis of unlinked SNPs, we identified 24 potential lineages within the five co-distributed complexes. Four of the five complexes exhibited contact zones between lineages at the Isthmus of La Paz, and all five did in the Vizcaíno Desert. To generate data for a species tree model, we subsampled two lizards from each of these 24 lineages for use with a target sequence capture (TSC) protocol. The resulting time-calibrated species tree shows that within each genus, the La Paz divergences are older than those across the Vizcaíno Desert. A full-likelihood Bayesian comparative phylogeographic approach was also used to test the simultaneous divergence hypothesis for the Isthmus of La Paz and Vizcaíno Desert, providing strong support for at least three independent divergence events at each contact zone, thereby rejecting the simultaneous divergence hypothesis. Finally, we demonstrate through expanded geographic sampling (n=142) and ddRAD that zebra-tailed lizards (*Callisaurus*), in which the most divergent lineages are endemic to the southern BCP, exhibit a clear pattern of Pleistocene range expansion from the northern BCP into the deserts of the western United States and mainland Mexico. The most deeply nested individuals in our tree occur at the northern, eastern, and southeastern range limits in temperate, subtropical, and tropical biomes, respectively. These results collectively highlight the importance of the BCP’s tectonic isolation as a driver not only of local or peninsular endemism, but potentially also as a contributing factor to lineage diversification more broadly in the region.

## INTRODUCTION

The Baja California Peninsula (BCP; Figure 1) is an ideal region to study biogeography across a range of spatial and temporal scales. Rugged, arid, and remote, with high topographic and climatic complexity, the BCP extends ∼1,400 km (12 degrees of latitude) from San Gorgonio Pass in southern California, USA to Cabo San Lucas in Baja California Sur, Mexico. This geological circumscription, also meaningful in a biogeographic sense, means that the northern limit of the peninsula is not at the Mexico-United States border, but rather is marked by the northernmost extension of the Gulf of California along the San Andreas Fault, which occurred in the Pliocene. Contributing to the complexity, the BCP has forty-five islands along its 3,300 km coastline, and its mountainous spine has several sky islands exceeding 3,000 m elevation in the north and 2,000 m in the south (Peralta-García et al., 2023). The BCP was transferred from the North American plate to the Pacific plate during the late Miocene and Pliocene as oblique extensional faulting, and later seafloor spreading, formed the Gulf of California, shearing the peninsula from the mainland and displacing it by nearly 500 km to the northwest (Bennett & Oskin, 2014; Lonsdale, 1989; Moore & Buffington, 1968; Stock & Hodges, 1989). Marine conditions initiated ∼7–10 Ma in the southern ‘proto-Gulf’ basin (Carreño, 1985, 1992; Dolby et al., 2015; Gastil, 1978; McCloy et al., 1988; Molina-Cruz, 1994), and by ∼6.3 Ma inundation had reached the northern Gulf, synchronous with plate boundary localization (Oskin & Stock, 2003a, 2003b). The archipelago in the Gulf of California is renowned as an UNESCO World Heritage Site and is protected by the Mexican government as a Biosphere Reserve (Cody et al., 2002), and the peninsula, along with the rest of the North American deserts, is considered a high biodiversity wilderness area at a global level (Mittermeier et al., 2003). For these reasons and more, the BCP and Gulf of California are emerging as a model system for historical biogeography, and squamate reptiles are frequent focal organisms due to their abundance, low vagility, and tendency towards local or regional endemism (Harrington et al., 2018; Leavitt et al., 2020; Pavón-Vázquez et al., 2024; Peralta-García et al., 2023; Sumarli et al., 2024; Valdivia-Carrillo et al., 2017).

**Figure 1.**
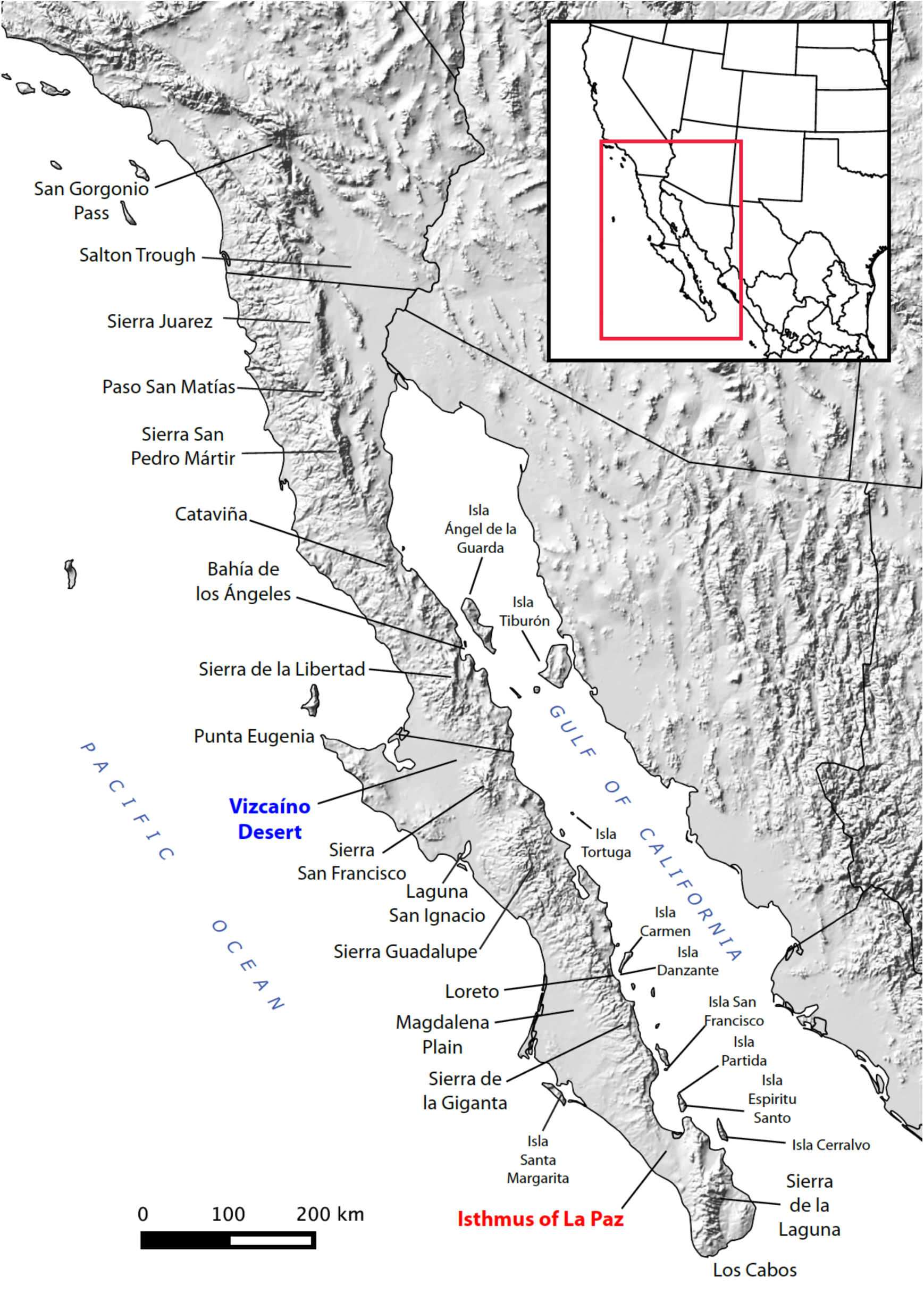
Map of Baja California Peninsula (BCP) with relevant geographic features mentioned in-text. Elevation data were obtained from the U.S. Geological Survey and Instituto Nacional de Estadística y Geografía (INEGI).

Spatially concordant breaks in mitochondrial DNA across terrestrial taxa have led some authors to treat the ancient BCP as a peninsular archipelago (Aguirre et al., 1999; Blair et al., 2009; Crews & Hedin, 2006; Lindell et al., 2005, 2006, 2008; Murphy, 1983; Murphy & Aguirre-León, 2002; Riddle et al., 2000; Upton & Murphy, 1997; Zink, 2002). This hypothesis posits that higher sea levels fragmented the peninsula in the mid-peninsula (Vizcaíno Desert) and/or the Isthmus of La Paz (Figure 1), two regions with low passes (∼400 m elevation) along the mountain ranges that separate the Gulf from the Pacific, although the timing and geological evidence is controversial (Dolby et al., 2015, 2024; Gardner et al., 2024). Similar phylogeographic patterns have been found in marine species (Lin et al., 2009; D. L. Peterson et al., 2013; Riginos, 2005). Other studies of peninsular taxa have emphasized the importance of glacial cycles of the Pleistocene epoch, causing fluctuations in sea level, temperature, and precipitation, and thus range contraction and expansion (Dolby et al., 2015; Garrick et al., 2009; Harrington et al., 2018; Valdivia-Carrillo et al., 2017).

Here, we use genome-wide sequence data to explore the diversification of peninsular phrynosomatid lizards in the context of the BCP’s late Cenozoic tectonic and climatic history. Phrynosomatids are ecologically diverse and widespread throughout the BCP and its islands, often showing spatially concordant phylogeographic or phenotypic breaks near the Vizcaíno Desert and Isthmus of La Paz. We address the following questions: 1) Was there shared divergence among taxa across the Vizcaíno Desert and the Isthmus of La Paz? 2) How many divergence events occurred, and which species were involved? 3) For *Callisaurus draconoides*, which ranges more widely in western North America, including the eastern side of the Gulf, how are continental lineages related to peninsular and insular lineages?

## MATERIALS & METHODS

### Field sampling

Details of the samples used in this study, including GPS coordinates, localities and museum voucher information, are provided in Supplemental Tables 1 & 2. Specimens and tissue samples were either collected by the authors in the United States and Mexico under the appropriate state and federal permits (see Acknowledgements) or borrowed from other individuals or museums. Prior to genomic work, liver and tail tip tissues were preserved in 96– 100% ethanol or RNAlater buffer and stored frozen at −20° C or −80° C.

### ddRAD library preparation

In order to identify the evolutionarily significant lineages for comparison (i.e., operational species under a species tree model), we first sampled 228 lizards (61 *Callisaurus*, 40 *Petrosaurus*, 47 *Urosaurus*, and 80 *Sceloporus*) spanning the BCP and eight islands (Supplemental Table 1). We collected genome-wide DNA sequence data using the double-digest Restriction-Associated-DNA (ddRAD) sequencing protocol (B. K. Peterson et al., 2012) and using the same enzymes and size selection window as Gottscho et al. (2017, 2024), described below. Hereafter we refer to this dataset as “BCP ddRAD”. We used the DNeasy Blood and Tissue Kit (Qiagen, Valencia, CA) with RNAse A to extract genomic DNA (gDNA). DNA concentrations were measured using a Qubit 2.0 Fluorometer (Life Technologies, Grand Island, NY) and assessed for high molecular weight using 1% agarose gel electrophoresis. The high-fidelity restriction enzymes *SbfI* and *MspI* (New England Biolabs, Ipswich, MA) were used to digest 200-500 ng of gDNA per sample. Digested samples were purified using Agencourt AMPure beads (Beckman Coulter, Danvers, MA) before attaching uniquely bar-coded adapters to each library with T4 Ligase (New England Biolabs, Ipswich, MA). After a second AMPure cleanup, samples were combined to create pools of eight uniquely barcoded libraries at equimolar concentrations. Fragments 415–515 bp long were selected using a Pippen Prep (Sage Science, Beverly, MA) with 2% gel cassettes. We used Phusion *Taq* Polymerase (New England Biolabs) for the Polymerase Chain Reaction (PCR) with a two-step cycle (98° C for 10 s, 72° C for 20 s) 12x, followed by a final extension step of 72° C for 10 min. An Agilent Bioanalyzer 2100 was used to ensure that the libraries were at the appropriate concentration and size distribution for sequencing. The final libraries (100 bp single-end reads) were sequenced on a HiSeq 2500 (Illumina, San Diego, CA) at the Institute of Integrative Genome Biology (University of California, Riverside).

### ddRAD bioinformatics

We used the Python pipeline pyRAD v3.0.6 for quality control, alignment, and genotype calling (Eaton, 2014). Raw reads were de-multiplexed by barcode, restriction cut sites and adapter sequences were trimmed, bases with Phred scores < 20 were replaced with ‘N’, and reads with >10 Ns were removed. Reads were clustered into loci for each individual separately using a clustering threshold of 0.85, then heterozygosity and error rates were jointly estimated for each individual. Consensus base calling was performed, retaining only loci with at least 10x coverage, and consensus sequences with >5 heterozygous sites were excluded to filter possible paralogs. Consensus sequences were clustered across individuals using a 0.85 clustering threshold, loci in which > 75% of samples had shared heterozygosity at a site were filtered, and alignments of both full loci and variable sites (individual SNPs) were generated for further analysis.

### Population structure and concatenated phylogeny

Admixture v1.23 (Alexander et al., 2009) was used to detect population structure. We used the cross-validation errors (CVE) to select the optimal K value ranging from 2 to 10, and used a custom R script to plot the results. A maximum-likelihood (ML) phylogeny of the concatenated data was constructed separately for each of the four clades with RAxML v8.2.7 (Stamatakis, 2014). The GTR+GAMMA substitution model using empirical base frequencies was implemented, using the BFGS algorithm to optimize GTR rate parameters. 1000 rapid bootstrap replicates were performed followed by a thorough ML search. Trees were initially evaluated unrooted or with midpoint rooting, and final root locations (Figures 2-5) are based on the species tree (Figure 6).

**Figure 2.**
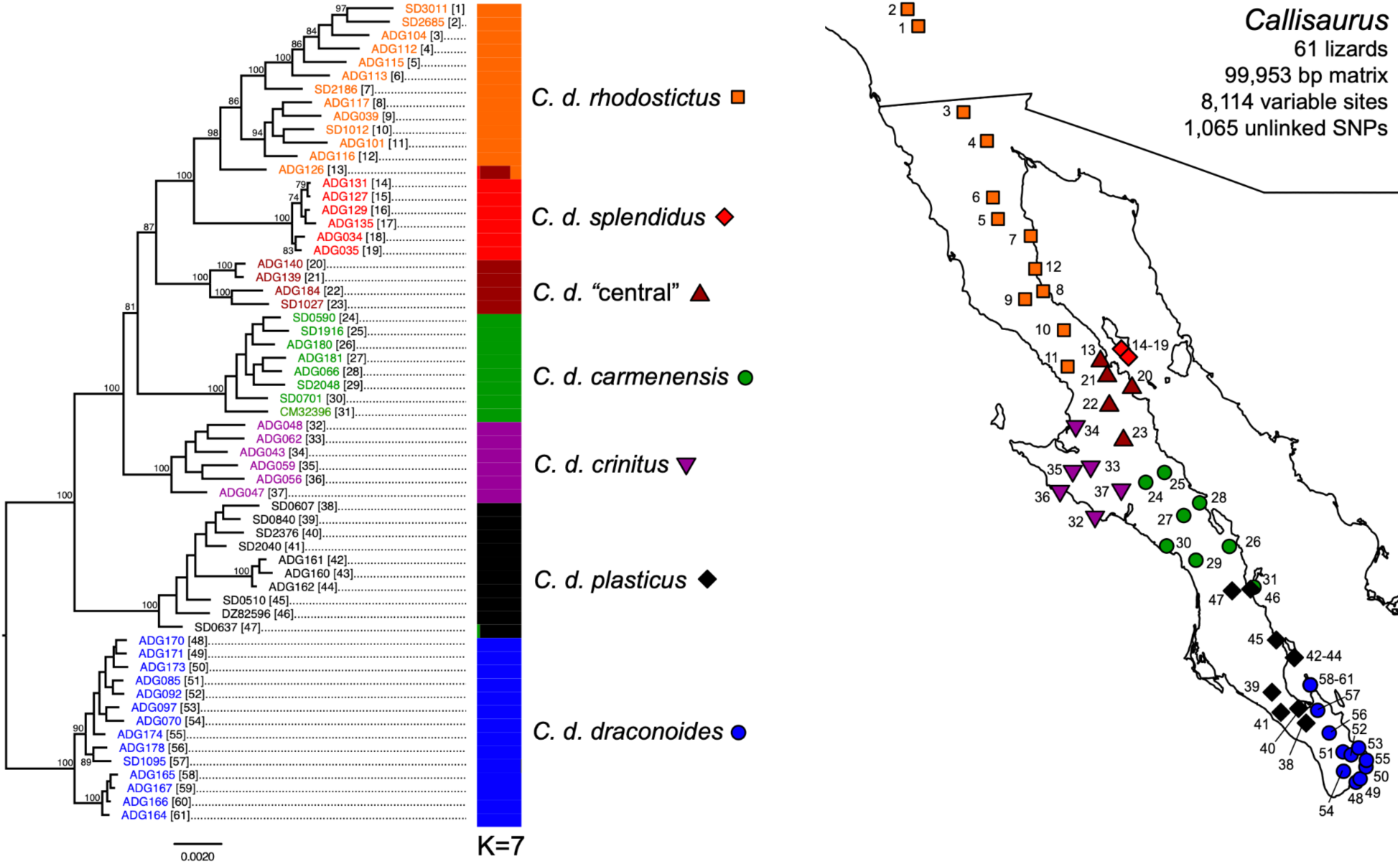
Maximum likelihood phylogram of concatenated ddRAD data (99,953 bp matrix, 8,115 variable sites) for peninsular *Callisaurus draconoides* (n=61). Clades with bootstrap support > 70% are shown on nodes, and numbers in brackets next to taxa labels (left) correspond to the numbers on the map (right). The middle bar plot represents the Admixture analysis of 1,065 unlinked SNPs under the optimal K=7.

### TSC data acquisition

We subsampled 48 individuals from the BCP ddRAD dataset (Supplemental Table 3), representing two individuals per each of 24 lineages inferred from the previous ddRAD analyses (see Results, Figures 2-5). Next, we used the MYbaits target enrichment kit (MYcroarray Inc., Ann Arbor, MI) to target 585 loci from these 48 lizards with 1,170 custom probes (two tiled 120 bp probes per locus overlapping by 60 bp) (Leaché et al., 2015). 541 of the targeted loci are ultra-conserved elements (Faircloth et al., 2012) with 99% sequence similarity to the *Anolis carolinensis* genome (Alföldi et al., 2011), and the remaining 44 genes were selected from the squamate Tree of Life project (Wiens et al., 2012). Hereafter, we refer to this as the target sequence capture (“TSC”) dataset, which was collected at the National Museum of Natural History, Laboratory of Analytical Biology. To prepare our DNA libraries with unique dual-index primers for each individual, we used the Kapa BioSystems Hyper Prep Kit with ¼ standard reaction volumes. We sheared gDNA extracts with a Qsonica DNA shearer by pipetting 250 ng into a shearing tube to a total volume of 110 µL, vortexing at 25% amplitude, 10-10 pulse, for 3 min. After end-repair and A-tailing, universal stubs were ligated. iTru i5 & i7 PCR primers were used in the PCR reaction (98° C for 45 seconds; 14 cycles of: 98° C for 15 seconds, 65° C for 30 seconds, and 72° C for 60 seconds; 72° C for 5 min). Hybridization, washing, library elution, and amplification followed the MYbaits Manual v3.01. Post-hybridization libraries were enriched using PCR with iTru adapter primers (98° C for 2 min; 14 cycles of: 98° C for 20 s, 60° C for 30 s, 72° C for 1 min; 72° C for 5 min) using KAPA HiFi HotStart RM polymerase. qPCR and a Bioanalyzer were used to assess library quality prior to sequencing. Libraries were pooled and sequenced on an Illumina NextSeq (150 x 150 paired-end reads) at the Institute of Integrative Genome Biology (University of California, Riverside, CA).

### TSC bioinformatics

The TSC FASTQ data were processed with a modified version of the phyluce v1.5 pipeline (Faircloth, 2016). Details of the steps and modifications are documented on github (https://github.com/SmithsonianWorkshops/Targeted_Enrichment). Trim Galore (https://github.com/FelixKrueger/TrimGalore), a wrapper around cutadapt and FastQC, was used to trim adapters and low quality reads. Trinity (https://github.com/trinityrnaseq/trinityrnaseq) was used to assemble data into contigs. We used LASTZ (https://github.com/lastz/last) to match contigs to the probe set and remove duplicates, i.e., loci that matched multiple contigs. Loci were extracted to FASTA format, with a separate FASTA file for each lizard. Loci were aligned with MAFFT and trimmed internally with Gblocks. NEXUS files for each locus were output at two levels of missing data (75% and 90% complete data matrices).

### Species tree

We first analyzed the 75% complete concatenated matrix with BEAST v2.4.5 (Bouckaert et al., 2014), primarily for quality control to ensure that two selected individuals per lineage indeed grouped together as sister taxa in the phylogeny. A coalescent species tree (24 species, 48 individuals) was inferred with StarBEAST2 v2.0.13.2 (Ogilvie et al., 2017) using a smaller dataset of 26 exons with >90% complete data (larger datasets faced convergence issues). We chose an uncorrelated lognormal relaxed clock, the Analytical Population Size Integration population model, and the HKY site model for all loci, with a gamma distribution on the kappa parameter. We set a calibrated Yule process species tree prior, with all clock rates to 1/X (uninformative) for each locus. We calibrated the molecular clock on both trees by setting a normally distributed prior on crown Phrynosomatidae of 55 Ma, sigma=4 (Leaché et al., 2015). All other priors were left as defaults. We ran three separate instances of 500 million Markov chain generations, sampling every 50,000 generations. Convergence of parameters was assessed with Tracer v1.6 (http://beast.bio.ed.ac.uk/Tracer). After discarding the first 20% of sampled generations from each run as burn-in, the maximum clade consensus tree was created with LogCombiner and TreeAnnotator (Bouckaert et al., 2014).

### Simultaneous divergence models

We used a full-likelihood Bayesian approach, ecoevolity (Oaks, 2019), to test the simultaneous divergence hypothesis for population pairs distributed across the Isthmus of La Paz and the Vizcaíno Desert. Ecoevolity treats each pair of populations as a species tree with two tips, and uses a Dirichlet process prior to estimate the number of shared divergence “events” (groups of contemporaneous individual divergences between population pairs) and the order of divergences. The assumptions of the model include constant population size, no migration, and that the relative mutation rates are similar among populations. Tests for shared divergence included four lineage pairs for the Isthmus of La Paz and seven for the Vizcaíno Desert. The Dirichlet process uses a concentration parameter to determine the prior probability for shared divergence events. We compared analyses using two extreme assumptions regarding this prior, including an “independent prior” that places approximately 50% of the probability on independent divergences for each pair, and a “shared prior” that places approximately 50% of the probability on all pairs sharing the same divergence time. Analyses were conducted using the BCP ddRAD data and the TSC data separately.

### *Callisaurus* range-wide ddRAD

To examine the relationships of peninsular *Callisaurus* to those distributed elsewhere in the North American deserts, we included an additional 79 specimens of *Callisaurus draconoides* and two specimens of *Holbrookia elegans* as outgroups (Supplemental Table 2). Most of these animals were collected in the states of California, Nevada, Utah, Arizona, New Mexico, Sonora, and Sinaloa, but we also included six new samples from isolated localities in Baja California Sur. Hereafter, we refer to this dataset, combined with the existing 61 peninsular *Callisaurus* from the BCP ddRAD dataset, as “*Callisaurus* range-wide ddRAD” (142 individuals).

We applied the same ddRAD methods and analytical approach as described above for peninsular *Callisaurus*, with slight differences described below. Libraries were pooled and sequenced on either an Illumina HiSeq 2500 (100 bp single-end reads) or a NextSeq (150 bp single-end reads) at the Institute of Integrative Genome Biology (University of California, Riverside, CA). Prior to running pyRAD, we trimmed the NextSeq reads to 100 bp. pyRAD v3.0.66 was run as described above for the BCP ddRAD data. We used RAxML v8.2.13 to produce a ML phylogram under the GTR+GAMMA substitution model on the concatenated ddRAD loci by executing 1000 rapid bootstrap inferences followed by a thorough ML search. Admixture v1.3.0 was used on the unlinked SNPs to assess population structure. Because the Eigenstrat (.geno) file generated by pyRAD is no longer supported in v1.3.0, to maintain continuity of methods, we converted the Structure (.str) file to PED format with PGDSpider v.3.0.0 (Lischer & Excoffier, 2012), then used PLINK v1.90b7.2 (Chang et al., 2015) to convert these to binary format that can be input by Admixture. We used the CVE to select the best-scoring K value (5–12) across three replicate runs with different random number seeds.

## RESULTS

### BCP ddRAD summary statistics

Raw ddRAD data characteristics from pyRAD are shown in Table 1 and Supplemental Table 4. After demultiplexing, we obtained a total of 255.6 million reads (mean of 1.12 million per lizard), of which 240.6 million passed the quality filter (mean of 1.06 million per lizard). Subsequent to clustering reads into putative loci, we obtained an average of 31,055 loci per lizard at an average sequencing depth of 35x; of these, an average of 5,389 loci passed the 10x coverage threshold for mean coverage of 198x. Concatenated data matrices of comparable size were generated for *Callisaurus* (99,953 bp matrix containing 8,114 variable sites), *Petrosaurus* (100,152 bp matrix, 6,663 variable sites), *Urosaurus* (96,389 bp matrix, 10,615 variable sites), and *Sceloporus* (97,476 bp matrix, 12,332 variable sites). For the population structure analyses, selecting one SNP per locus and filtering for biallelic SNPs resulted in 1,065 unlinked SNPs for *Callisaurus*, 1,046 for *Petrosaurus*, 1,024 for *Urosaurus*, and 1,039 for *Sceloporus*.

**Table 1.** BCP ddRAD data summary. Reads Passed = mean number of raw reads that passed initial quality filter after demultiplexing; Clusters = mean number of clusters assembled *de novo* that passed 10x coverage threshold; Coverage = Mean depth of coverage (x) for clusters that passed 10x coverage threshold; F2 Loci = mean number of clusters that passed coverage threshold and paralog filter; Het = mean heterozygosity (percent); Final Loci = mean number of loci recovered for each taxon in the final dataset after alignment and final filtering; n = number of individuals in the final dataset (sum).

|  | Reads Passed | Clusters | Coverage | F2 Loci | Het | Final Loci | n |
| --- | --- | --- | --- | --- | --- | --- | --- |
| *Callisaurus* | 1,130,138 | 5,655 | 195.8 | 5,328 | 0.36% | 986.2 | 61 |
| *Petrosaurus* | 1,047,981 | 6,520 | 200.1 | 6,030 | 0.26% | 828.7 | 40 |
| *Urosaurus* | 1,147,850 | 3,641 | 241.0 | 3,426 | 0.45% | 767.9 | 47 |
| *Sceloporus* | 947,430 | 5,648 | 174.6 | 5,274 | 0.40% | 777.2 | 80 |
| All Taxa | 1,055,268 | 5,389 | 198.4 | 5,040 | 0.37% | 840.2 | 228 |

### *Callisaurus* phylogeography and population structure

The results of the Admixture analysis found that K=7 was the best supported model (Table 2), and these seven clusters nearly perfectly match seven well-supported clades (bootstrap values ≥0.70) from the RAxML analysis (Figure 2). Four of the seven closely match previously recognized subspecies, treated as “pattern classes” by Grismer (2002): 1) *C. d. rhodostictus* (Cope, 1896) from the northernmost BCP, north of Bahía de los Ángeles, 2) *C. d. splendidus* (Dickerson, 1919) from Isla Ángel de la Guarda, 3) *C. d. crinitus* (Cope, 1896) of the western Vizcaíno Desert, and 4) *C. d. draconoides* (Blainville, 1835) from the southernmost Cape Region (including Isla Espiritu Santo). The clade and cluster 5) *C. d. carmenensis* (Dickerson, 1919), as newly delimited here, ranges from south of the Sierra de San Francisco to Isla Carmen, and that 6) *C. d. plasticus* (Dickerson, 1919), which had been subsumed within *C. d. carmenensis* (Grismer, 2002; Van Denburgh, 1922), is distributed from Isla Danzante to the Isthmus of La Paz (including Isla San Francisco).The *C. d.* “central” population 7) does not match any previously recognized subspecies and has a small distribution along the central Gulf coast, east of the Vizcaíno Desert and north of the Sierra de San Francisco. Its range falls within an area previously depicted as an intergrade zone (Grismer, 2002). One individual (Field No. ADG126) from just west of Bahía de los Ángeles, which groups with *C. d. rhodostictus* in the tree, exhibits admixture between *C. d. rhodostictus* and *C. d.* “central” (Figure 2). This individual also has the highest proportion of heterozygous sites of any *Callisaurus* (Supplemental Table 4). Unlike lizards from Isla Ángel de la Guarda, the insular populations on Isla Espiritu Santo, Isla San Francisco, Isla Danzante, and Isla Carmen were not recognized as distinct clusters under K=7.

**Table 2.**
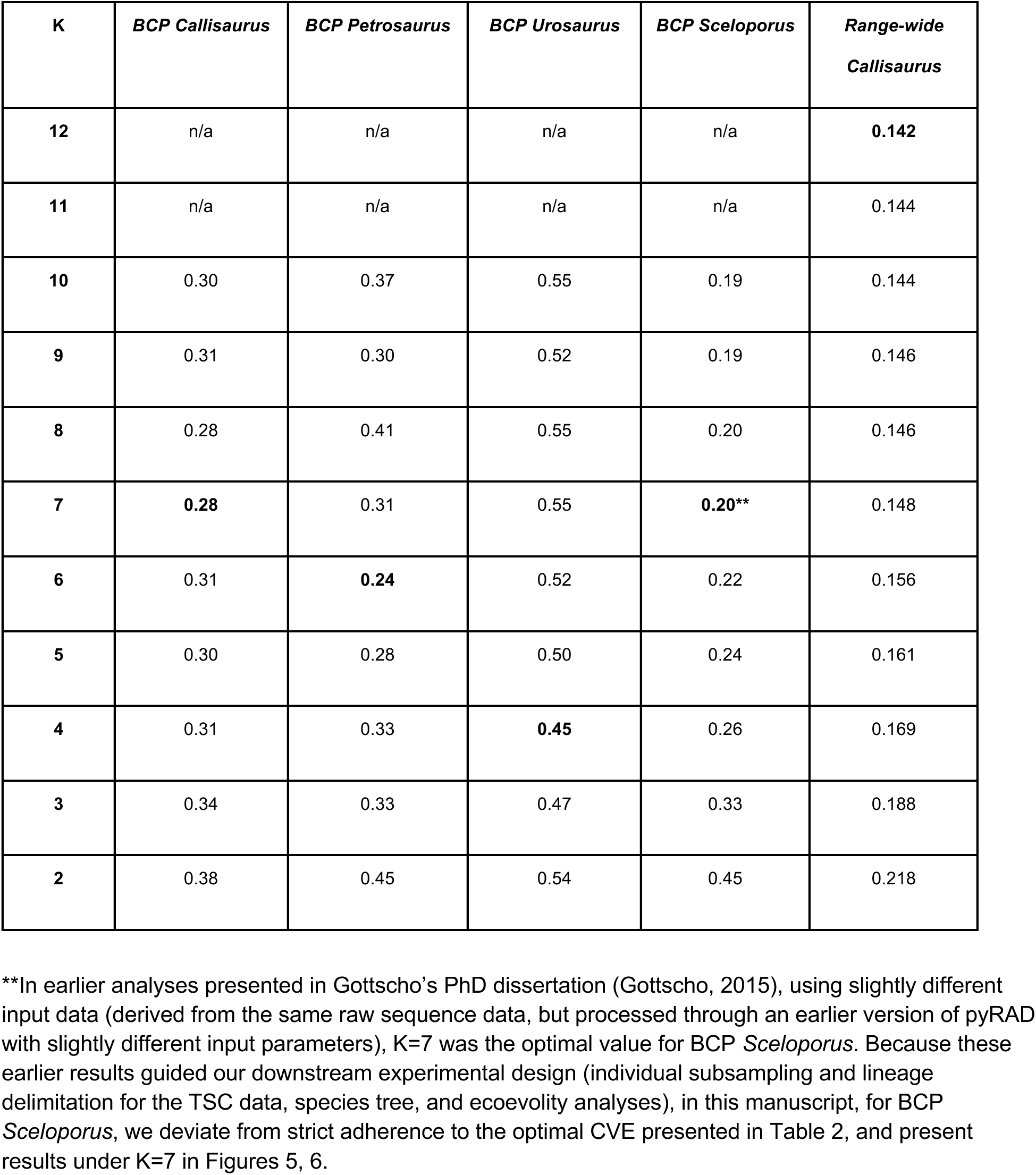
Cross-validation error (CVE) results from Admixture (BCP ddRAD). Boldface values indicate selected scores for visualization (Figures 2–5, 9) and for identifying lineages in the species tree (Figure 6). In most cases, these were the optimal values, except for BCP *Sceloporus* where K=7 was chosen (see footnote **). Values K=2–10 were tested for the BCP ddRAD data in Admixture v1.23, and K=2–12 for the range-wide *Callisaurus* data in Admixture v1.3.0.

| K | *BCP Callisaurus* | *BCP Petrosaurus* | *BCP Urosaurus* | *BCP Sceloporus* | *Range-wide Callisaurus* |
| --- | --- | --- | --- | --- | --- |
| 12 | n/a | n/a | n/a | n/a | **0.142** |
| 11 | n/a | n/a | n/a | n/a | 0.144 |
| 10 | 0.30 | 0.37 | 0.55 | 0.19 | 0.144 |
| 9 | 0.31 | 0.30 | 0.52 | 0.19 | 0.146 |
| 8 | 0.28 | 0.41 | 0.55 | 0.20 | 0.146 |
| 7 | **0.28** | 0.31 | 0.55 | **0.20\*\*** | 0.148 |
| 6 | 0.31 | **0.24** | 0.52 | 0.22 | 0.156 |
| 5 | 0.30 | 0.28 | 0.50 | 0.24 | 0.161 |
| 4 | 0.31 | 0.33 | **0.45** | 0.26 | 0.169 |
| 3 | 0.34 | 0.33 | 0.47 | 0.33 | 0.188 |
| 2 | 0.38 | 0.45 | 0.54 | 0.45 | 0.218 |

### *Petrosaurus* phylogeography and population structure

The RAxML tree (Figure 3) for *Petrosaurus* revealed six strongly supported clades that were also identified as clusters by Admixture under the best fit model (K=6; Table 2): 1) a northern clade of *P. mearnsi* (Stejneger, 1894) that ranges from San Gorgonio Pass to the east slope of the Sierra San Pedro Martír, 2) a southern clade consisting of *P. mearnsi* ranging from Cataviña to Bahía de los Ángeles and *P. slevini* (Van Denburgh, 1922) from Isla Ángel de la Guarda (hereafter, the entire southern clade is referred to as *P. “slevini”*), 3) a northern clade of *P. repens* (Van Denburgh, 1895) found northwest of Bahía de los Ángeles, 4) a central clade of *P. repens* found in the Sierra de San Francisco, Sierra Guadalupe, and the northern Sierra de la Giganta, 5) a southern clade of *P. repens* ranging from the southern Sierra de la Giganta to the Isthmus of La Paz, and 6) *P. thalassinus* (Cope, 1863) from the Cape Region (Isla Partida to Los Cabos).

**Figure 3.**
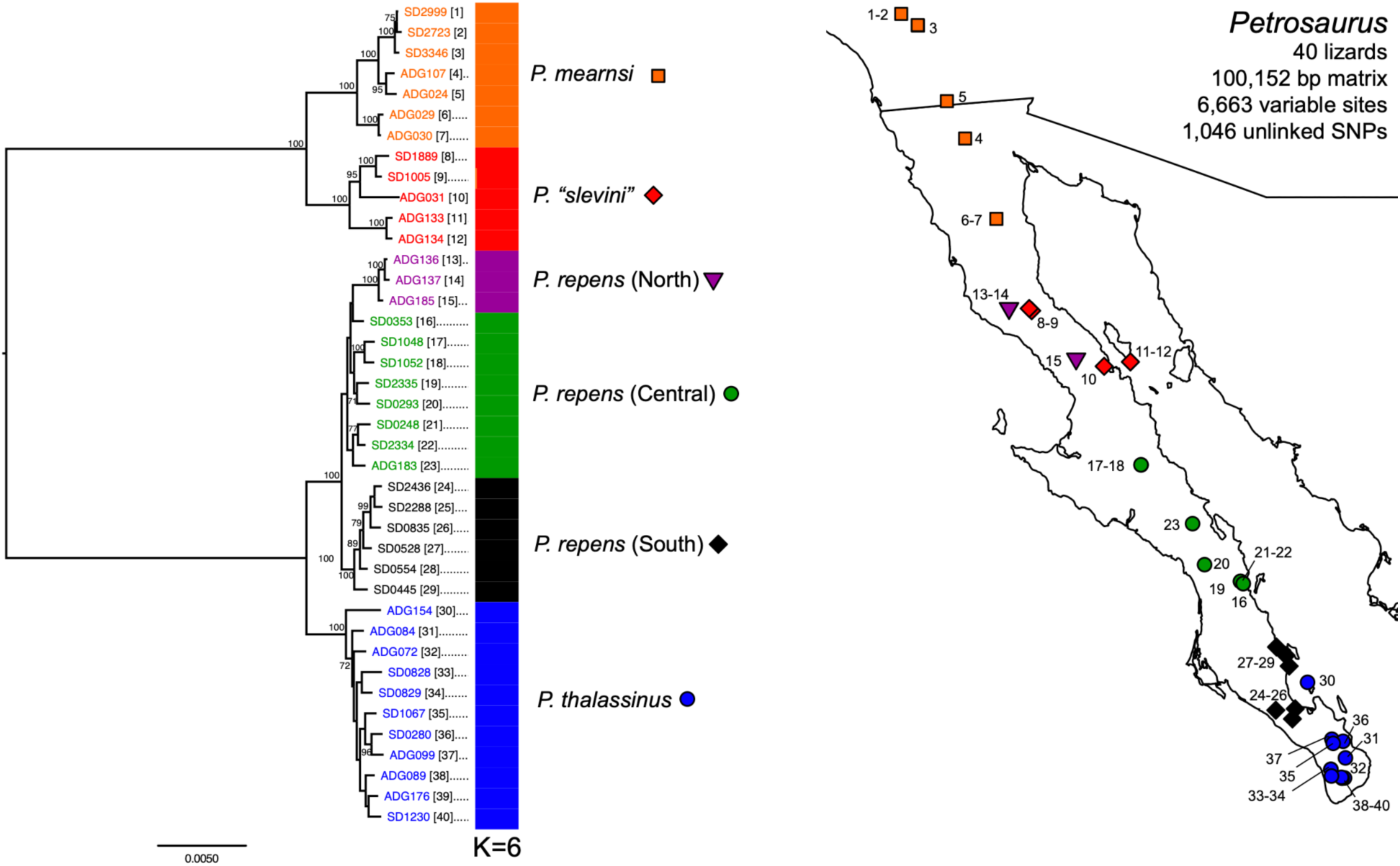
Maximum likelihood phylogram of concatenated ddRAD data (100,152 bp matrix, 6,663 variable sites) for *Petrosaurus* (n=40). Clades with bootstrap support > 70% are shown on nodes, and numbers in brackets next to taxa labels (left) correspond to the numbers on the map (right). The middle bar plot represents the Admixture analysis of 1,046 unlinked SNPs under the optimal K=6.

### *Urosaurus* phylogeography and population structure

The Admixture analysis identified K=4 as the best-fit model; three of the four clusters are congruent with clades in the RAxML tree, and one cluster matches a paraphyletic group (Figure 4). 1) One cluster corresponds to *Urosaurus lahtelai* (Rau & Loomis, 1977), restricted to the vicinity of Cataviña, which is sister to all remaining individuals in the tree. The second cluster 2) consists of *U. nigricauda* (Cope, 1864) ranging from Isla Santa Margarita on the Pacific coast to Los Cabos, which is a strongly supported clade in RAxML sister to *U. microscutatus*, a previously described species (Van Denburgh, 1894) that was later subsumed within *U. nigricauda* (Aguirre et al., 1999). The third cluster 3) is composed of *U. microscutatus* along the central Gulf coast, south of the Sierra de San Francisco to the Sierra de la Giganta, which is paraphyletic with respect to the fourth cluster 4), which groups monophyletic northern *U. microscutatus* from the Sierra de la Libertad to southern California. Five geographically intermediate *U. microscutatus* from the Sierra de San Francisco show limited admixture between the third and fourth clusters (Figure 4). Within *U. microscutatus*, there is a clear pattern in the RAxML tree whereby northwestern individuals are more deeply nested than southeastern ones.

**Figure 4.**
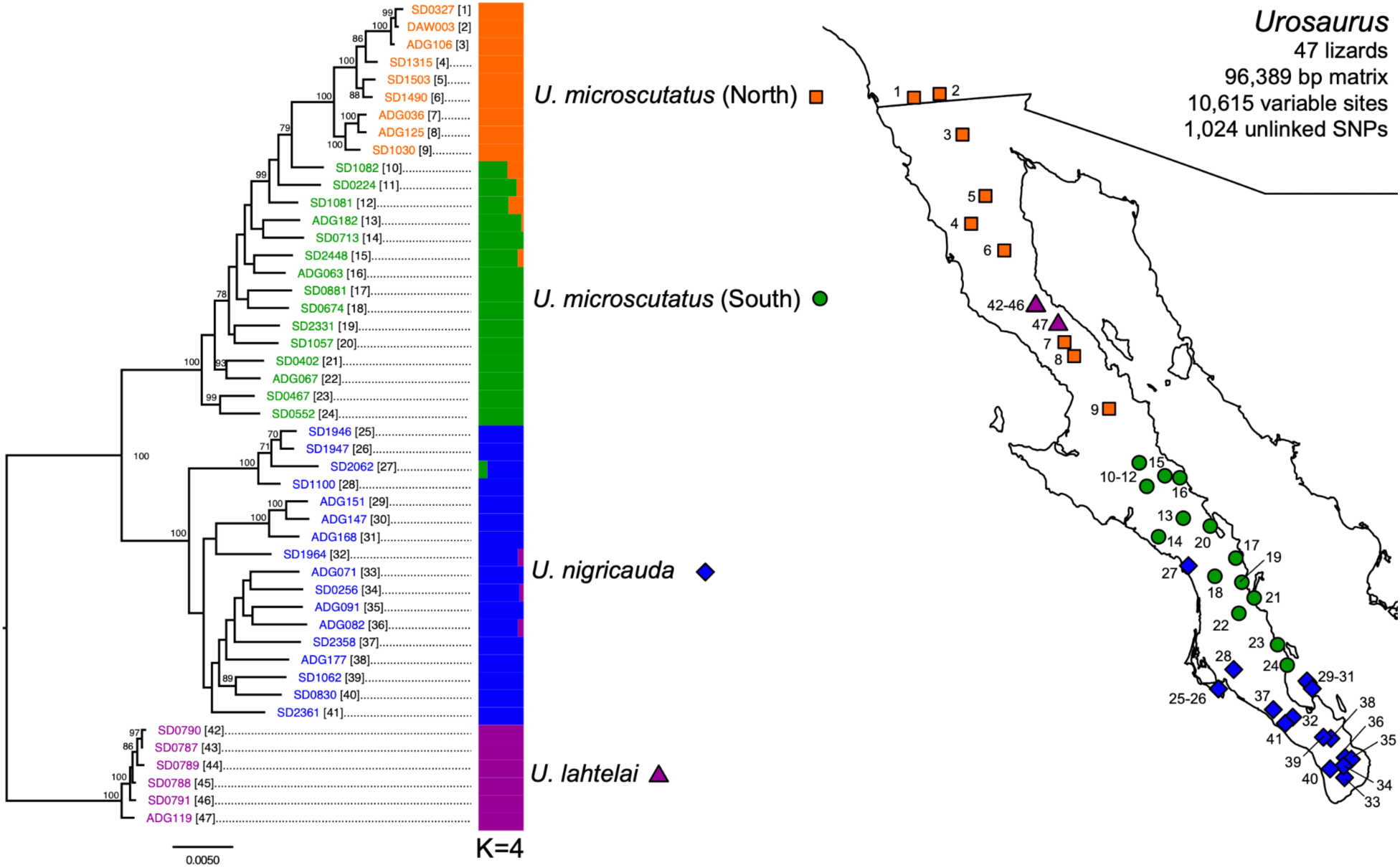
Maximum likelihood phylogram of concatenated ddRAD data (96,389 bp matrix, 10,615 variable sites) for peninsular *Urosaurus* (n=47). Clades with bootstrap support > 70% are shown on nodes, and numbers in brackets next to taxa labels (left) correspond to the numbers on the map (right). The middle bar plot represents the Admixture analysis of 1,024 unlinked SNPs under the optimal K=4.

### *Sceloporus* phylogeography and population structure

The clusters identified in Admixture (K=7, Table 2) match seven strongly supported clades (Figure 5) within two co-distributed complexes. Within the *S. orcutti* complex, we found 1) a northern clade of *S. orcutti* that ranges from San Gorgonio Pass to Cataviña, 2) a southern clade of *S. orcutti* (Stejneger, 1893) that ranges from the Sierra de San Francisco to the Sierra de la Giganta (with a substantial sampling gap between this and the northern clade), 3) *S. licki* (Van Denburgh, 1895) from the Cape Region (Sierra de la Laguna), and 4) *S. hunsakeri* (Hall & Smith, 1979) also from the Cape Region (Isla Partida to Los Cabos). In the sister *S. magister* complex, we found 5) a clade of *S. magister* (Hallowell, 1854) in the lower Colorado River valley sister to 6) a northern clade that corresponds closely to the previously recognized *S. zosteromus rufidorsum* (Grismer, 2002; Yarrow, 1882) ranging from Paso San Matías to the Vizcaíno Desert, and 7) a southern clade that corresponds to the combined previously recognized subspecies *S. z. monserratensis* and *S. z. zosteromus* (Cope, 1863; Grismer, 2002), and therefore takes the latter name, extending from the southern Vizcaíno Desert to Los Cabos.

**Figure 5.**
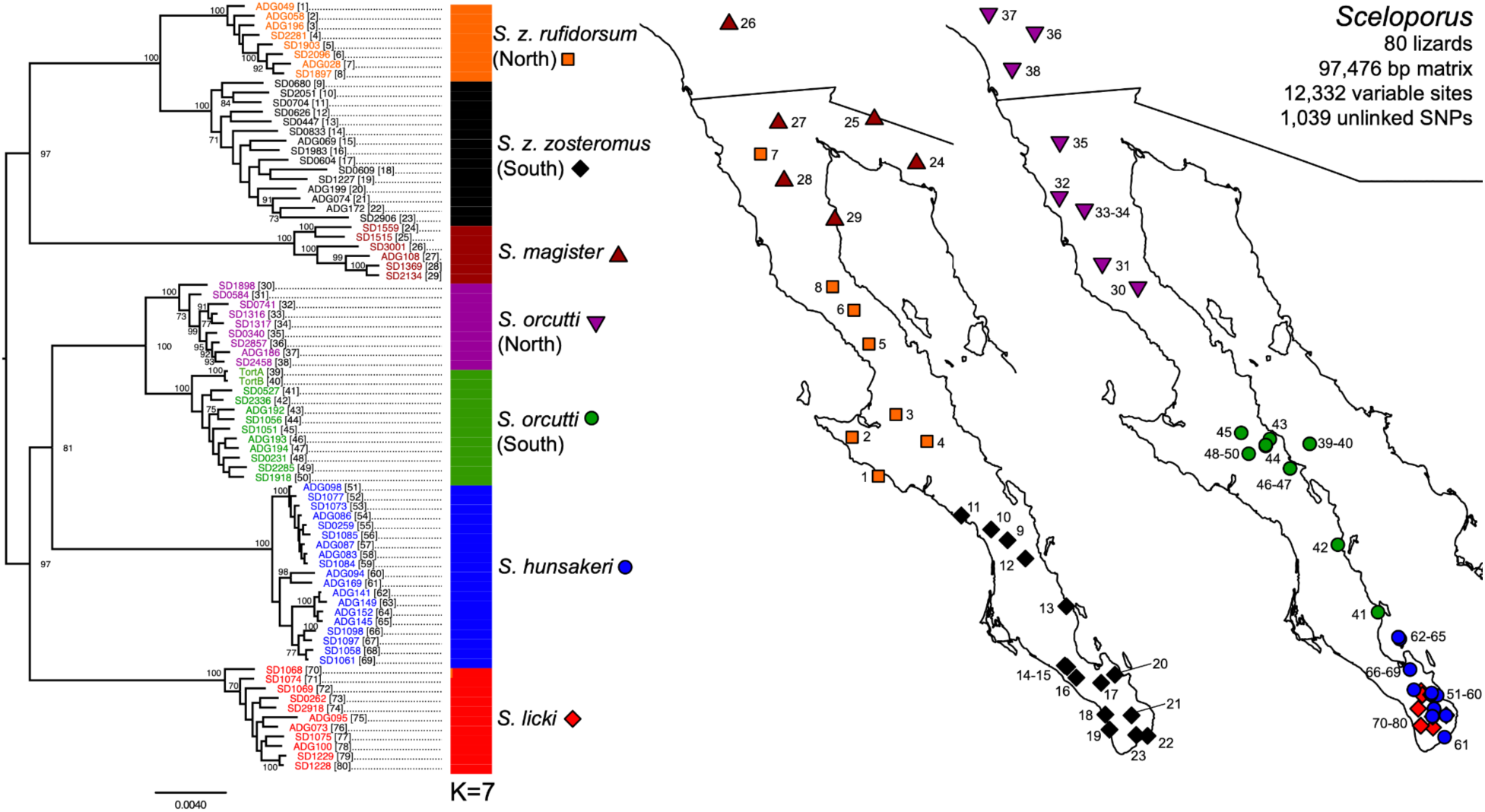
Maximum likelihood phylogram of concatenated ddRAD data (97,476 bp matrix, 12,332 variable sites) for peninsular *Sceloporus* included in this study (n=80). Clades with bootstrap support > 70% are shown on nodes, and numbers in brackets next to taxa labels (left) correspond to the numbers on the maps (right). The middle bar plot represents the Admixture analysis of 1,039 unlinked SNPs under K=7 (see footnote in Table 2).

### TSC summary statistics

TSC data characteristics are summarized in Table 3 and presented in detail in Supplemental Table 5. After demultiplexing, we obtained a total of 36.6 million reads (mean of 762,976 reads per lizard), at a mean contig coverage of 9x subsequent to assembly. A mean of 509 loci (87% of the targets) with a mean length of 737 bp were captured per lizard at a mean locus coverage of 34x.

**Table 3.** TSC data summary. Reads Passed = mean reads passed filter after demultiplexing (step 1); Contigs = mean number of Contigs from Trinity assembly; Contig Length = mean length of contigs assembled under Trinity; Contig Coverage = mean coverage of contigs assembled by Trinity; Loci Captured = mean number of targeted loci recovered (out of 585 total); Locus Length = mean length of targeted loci; Locus coverage = mean coverage of targeted loci; n = number of individual lizards per dataset (sum).

|  | Reads<br>Passed | Contigs | Contig<br>Length | Contig<br>Coverage | Loci<br>Captured | Locus<br>Length | Locus<br>Coverage | n |
| --- | --- | --- | --- | --- | --- | --- | --- | --- |
| *Callisaurus* | 806,295 | 14,036 | 313.5 | 8.6 | 506 | 742.5 | 39.0 | 14 |
| *Petrosaurus* | 665,354 | 13,401 | 322.3 | 8.7 | 510 | 809.2 | 40.5 | 12 |
| *Urosaurus* | 786,413 | 9,783 | 323.0 | 9.5 | 511 | 699.4 | 27.2 | 8 |
| *Sceloporus* | 788,659 | 9,689 | 324.1 | 9.4 | 510 | 694.2 | 26.5 | 14 |
| All Taxa | 762,976 | 11,885 | 320.5 | 9.0 | 509 | 737.0 | 33.6 | 48 |

### Species tree

The BEAST tree for the 75% complete concatenated matrix (549 loci, 48,023 bp) is shown in Supplemental Figure 1. This tree, which makes no assumptions about assignment of individual lizards to species, was primarily used as quality control to ensure that the two individuals sampled for each lineage are indeed sister to each other. The coalescent species tree (Figure 6) of 26 exons estimated with StarBEAST2 is our preferred tree for interpreting divergence dates and topologies within each genus. Both trees assume a crown age of 55 Ma (Leaché et al., 2015). The deepest nodes in the species tree (Figure 6) are consistent with Figure 1 in Leaché et al. (2015): *Callisaurus* diverged first, followed by *Petrosaurus*, which is sister to a clade containing *Urosaurus* and *Sceloporus*. This result was also seen in the concatenated tree (Supplemental Figure 1). All reported ages in this section are crown-group ages, geological epoch descriptions reflect the 95% highest posterior density (95% HPD), and node support was assessed with posterior probabilities (PP).

**Figure 6.**
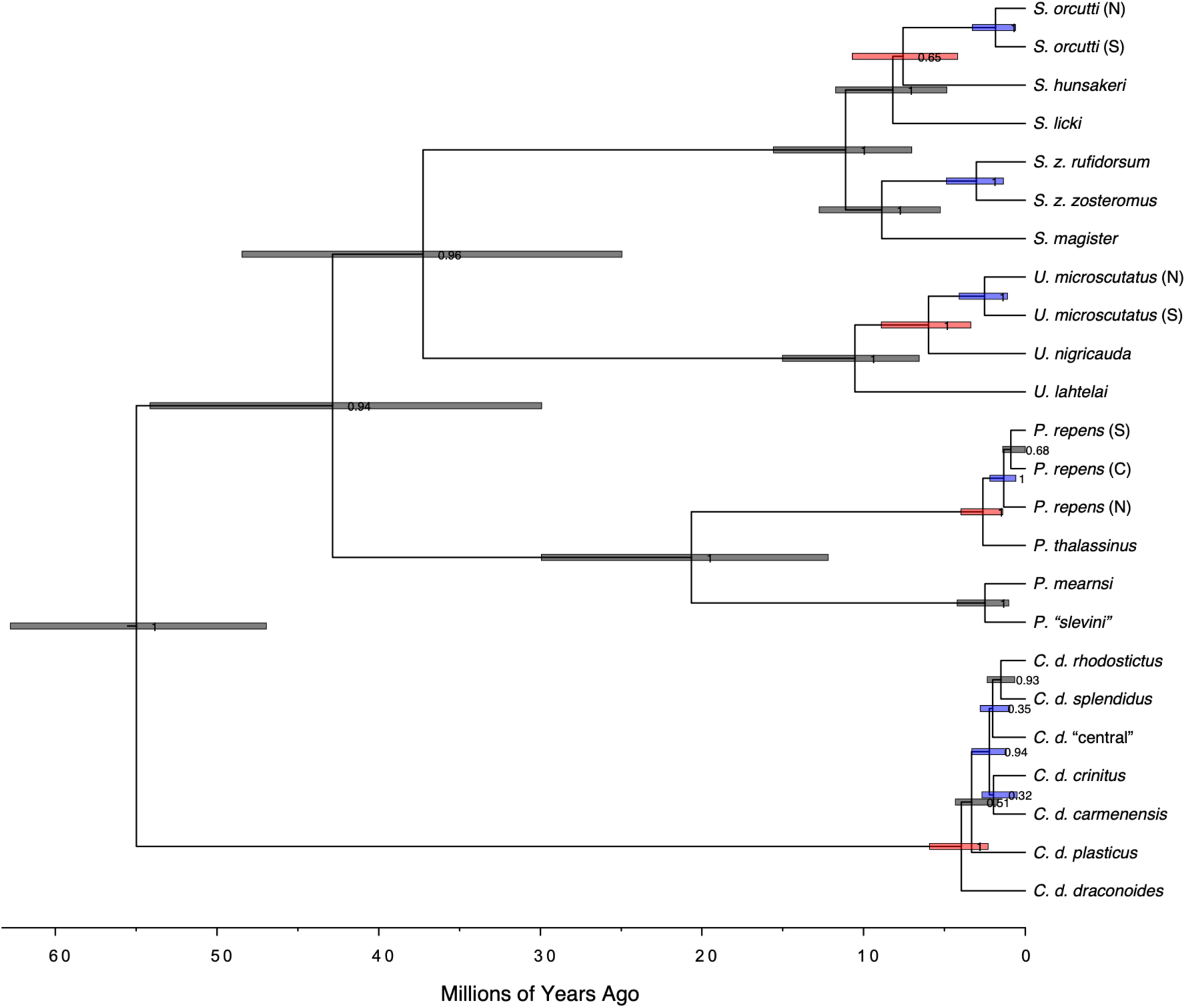
Time-calibrated species tree for BCP phrynosomatids based on 26 exons generated with StarBEAST2. Bars on nodes represent 95% highest posterior densities (HPDs) for divergence times, and numbers on nodes represent posterior probabilities (PP) for node support. Blue bars indicate that the lineages diverged across the Vizcaíno Desert break, and red bars indicate the Isthmus of La Paz break.

The age of peninsular *Callisaurus* was estimated to be 3.96 Ma (95% HPD 2.3–5.92 Ma), spanning the late Miocene to the early Pleistocene. The deepest split corresponds to the Isthmus of La Paz, between *C. d. plasticus* and *C. d. draconoides*, but the placement of *C. d. plasticus* is not strongly supported (PP=0.51). The monophyly of all other peninsular *Callisaurus* is strongly supported (PP=0.94), with a late Pliocene / early Pleistocene age of 2.19 Ma (95% HPD 1.19–3.31 Ma). Multiple lineages distributed north of (*C. d.* “central”), south of (*C. d. carmenensis*), or within (*C. d. crinitus*) the Vizcaíno Desert, diverged in rapid succession after this node. The placement of the *C. d.* “central” population was sister to a clade of *C. d. rhodostictus* and *C. d. splendidus*, although its placement was poorly supported. By contrast, the sister relationship of *C. d. rhodostictus* and *C. d. splendidus* was strongly supported (PP=0.93). The estimated date of divergence between *C. d. splendidus* and *C. d. rhodostictus* (1.48 Ma; 95% HPD 0.66–2.36 Ma) falls in the Pleistocene, and although this date does not bear on the questions regarding the Isthmus of La Paz and Vizcaíno Desert, it provides important context for interpreting the range-wide *Callisaurus* ddRAD dataset later in this article.

The crown age for *Petrosaurus* was estimated at 20.64 Ma (95% HPD 12.19–29.94 Ma), and although this clade is currently endemic to the BCP as defined earlier, this date is more than twice as old as the Gulf of California. Within *Petrosaurus*, there are two strongly supported clades (PP=1.0 in both cases). The first clade consists of northern *P. mearnsi,* in turn sister to a clade containing *P. “slevini”*, with a Plio-Pleistocene age of 2.49 Ma (95% HPD 1.02–4.22 Ma). The second clade contains the sister taxa *P. repens* and *P. thalassinus*, also of Plio-Pleistocene age (2.49 Ma; 95% HPD 1.39–3.98 Ma), and this divergence is across the Isthmus of La Paz. In *Petrosaurus*, the Vizcaíno Desert break occurs between northern and central *P. repens* (PP=1.0), and these lineages are estimated to have diverged in the Pleistocene (1.33 Ma; 95% HPD 0.59–2.19 Ma).

All nodes within the *Urosaurus* clade were strongly supported (PP=1.0). The age of peninsular *Urosaurus* included in the tree is 10.54 Ma (95% HPD 6.56–15.01 Ma), pre-dating the BCP, and corresponding to the divergence between *U. lahtelai* and the *U. nigricauda* complex. Within the *U. nigricauda* complex, the deepest divergence is between *U. nigricauda* and *U. microscutatus* across the Isthmus of La Paz in the late Miocene/Pliocene (5.98 Ma; 95% HPD 3.37 – 8.9 Ma). The divergence between northern and southern *U. microscutatus* across the Vizcaíno Desert was inferred to be of Plio-Pleistocene age (2.52 Ma; 95% HPD 1.1–4.09 Ma).

The crown age for the *Sceloporus* sampled was entirely Miocene (11.11 Ma; 95% HPD 7.03–15.57 Ma). One clade, the *S. magister* complex, with strongly supported monophyly (PP=1.0), has a late to mid-Miocene age of 10.54 Ma (95% HPD 5.26–12.74 Ma), which corresponds to the split between peninsular *S. zosteromus* and mainland *S. magister*. The split between *S. z. rufidorsum* and *S. z. zosteromus* across the Vizcaíno Desert was dated to the Plio-Pleistocene (3.03 Ma; 95% HPD 1.35–4.88 Ma). Sister to the *S. magister* complex is the *S. orcutti* complex, strongly supported as monophyletic (PP=1.0). *Sceloporus licki*, a Cape Region endemic, is sister to a clade containing *S. hunsakeri,* also a cape endemic, and *S. orcutti*. This latter clade was weakly supported (PP=0.65), but the divergence between *S. hunsakeri* and *S. orcutti* occurs across the Isthmus of La Paz and was dated to the late Miocene / early Pliocene (7.31 Ma; 95% HPD 4.18–10.7 Ma). The divergence between northern and southern *S. orcutti*, across the Vizcaíno Desert, was estimated to have occurred in the Plio-Pleistocene (1.94 Ma; 95% HPD 0.61–3.27 Ma).

### Comparative phylogeographic models

The ecoevolity analyses rejected the hypothesis of shared divergence at the Isthmus of La Paz and the Vizcaíno Desert. Instead, at least three independent divergence episodes are supported at each biogeographic barrier (Figure 7). For the Isthmus of La Paz, comparisons of analyses using priors that favored independent divergence versus shared divergence had little influence on the PP distributions for the number of events. For example, the ddRAD and TSC data both support three separate divergence events under each of the priors. For the Vizcaíno Desert, the dataset and the prior had an influence on the PP distributions for the number of events. First, the estimated number of events is lower for the TSC data (three) compared to the ddRAD data (four–five), and this pattern holds under both priors. Second, the independent prior tends to favor separate events more than the shared prior.

**Figure 7.**
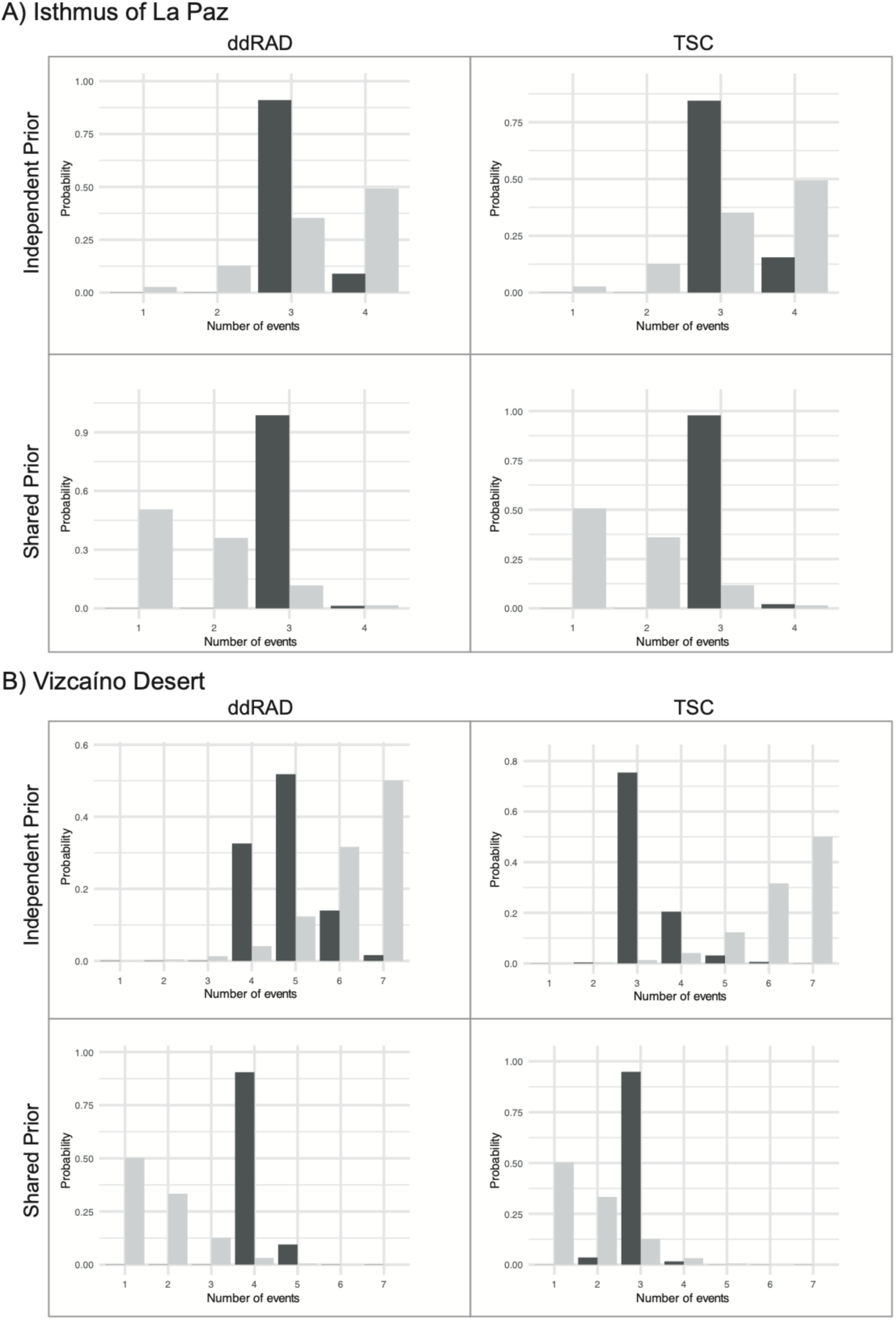
Comparative phylogeographic analysis results for the number of divergence events estimated for the Isthmus of La Paz (A) and Vizcaíno Desert (B) using ddRAD and TSC data. The analyses conducted assuming independent and shared priors are shown (priors are in light grey; posteriors in black).

The ordering of the four divergence events for the Isthmus of La Paz suggests that the oldest divergence is for *Sceloporus,* the most recent is for *Petrosaurus*, and *Urosaurus* is intermediate between the two (Figure 8). The placement of the *Callisaurus* divergence is uncertain, as it groups with *Petrosaurus* using the ddRAD data or with *Urosaurus* using the TSC data (Figure 8). The Vizcaíno Desert analyses included seven population pairs, and the TSC data support three divergence events with *Petrosaurus* occurring most recently, followed by *Urosaurus,* and then the remaining five population/species pairs grouped together. The ddRAD data support a different divergence history for the Vizcaíno Desert. Specifically, the divergence between *S. z. rufidorsum* and *S. z. zosteromus* is supported as an independent event that is also the oldest. Also, one of the pairs of divergences among the three *Callisaurus* lineages is inferred to be younger (though not as young as *Petrosaurus*) than the four population/species pairs that make up a common divergence event, which now includes *Urosaurus*.

**Figure 8.**
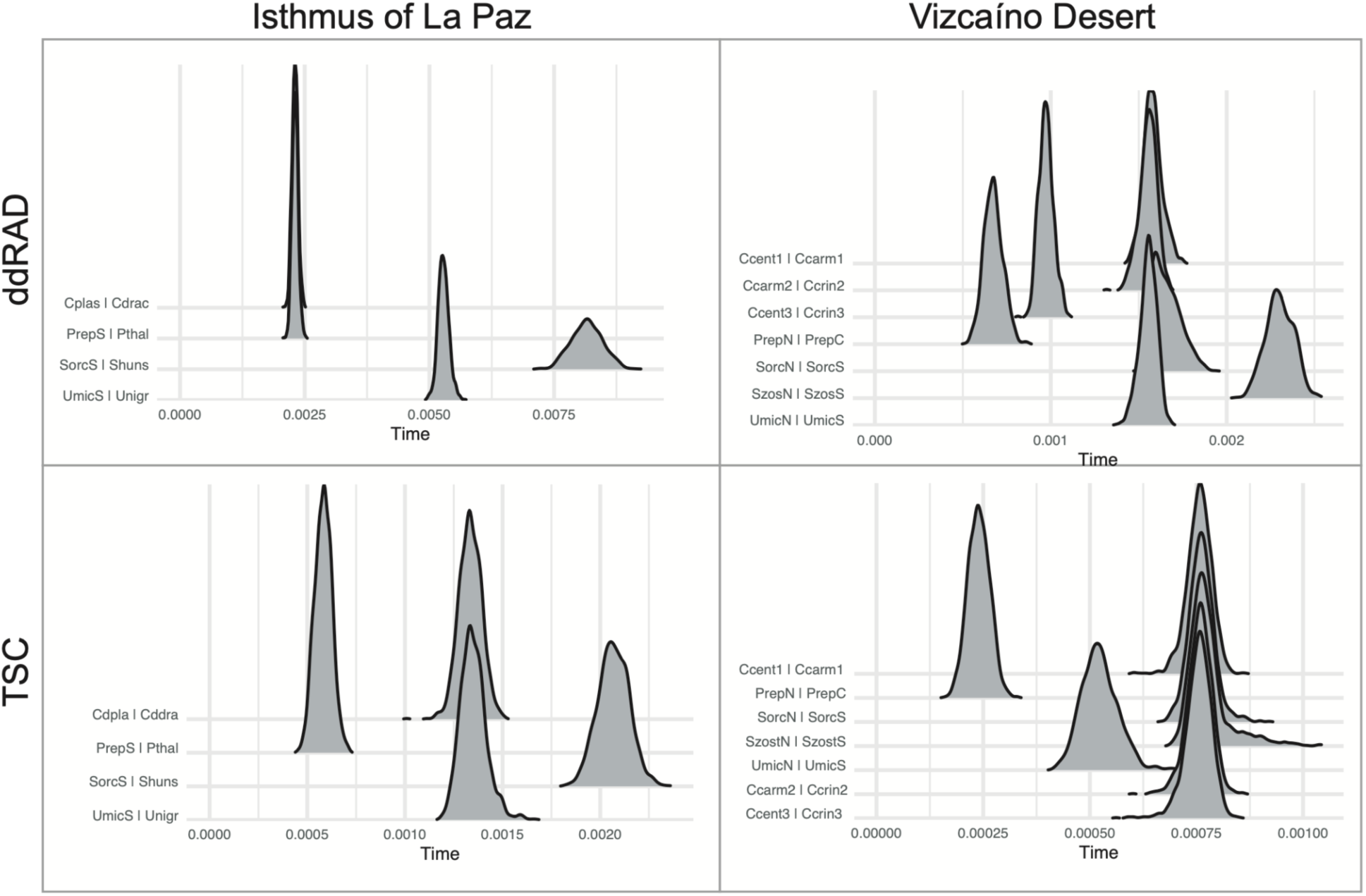
Posterior probabilities of divergence times across lineage pairs separated by the Isthmus of La Paz and the Vizcaíno Desert estimated using ecoevolity (time is expressed as substitutions/site on the x-axis) with ddRAD and TSC data. Results are shown for the independent prior.

### Callisaurus range-wide ddRAD

Figure 9 and Supplemental Figure 2 depict the RAxML tree for the concatenated *Callisaurus* range-wide ddRAD dataset (n=140) rooted with an outgroup (n=2), alongside the Admixture cluster assignments based on unlinked SNPs for the ingroup (n=140). The root location for the former differs from the species tree (Figure 6) in that the deepest divergence within *Callisaurus* is between a clade of *C. d. draconoides* + *C. d. plasticus* and all remaining *Callisaurus* (this result was also seen in the clock-rooted concatenated TSC tree, Supplemental Figure 1). The *C. d. draconoides* clade also has expanded sampling in the range-wide tree (Figure 9), with samples from the deep-water Isla Cerralvo, lying offshore the Cape Region, now falling within this clade. Otherwise, the broad phylogeographic patterns within the BCP are very similar to the peninsula-only tree (Figure 2). In the Admixture analysis, K=12 was the best-scoring model by CVE, and four of the clusters are peninsular and correspond to those found earlier for 1) *C. d. draconoides*, 2) *C. d. plasticus*, 3) *C. d. carmenensis*, and 4) *C. d. splendidus*. Admixture grouped 5) the *C. d.* “central” population with *C. d. crinitus*, and split peninsular *C. d. rhodostictus* into 6) southern (Central Desert) and 7) northern (Salton Trough) populations. In the phylogeny, *C. d. splendidus* of Isla Ángel de la Guarda was found to be monophyletic and sister to all other *C. draconoides* north of Bahía de los Ángeles and beyond—including all of the additional sampled *C. draconoides* in the United States and mainland Mexico—although not strongly supported. The addition of mainland samples revealed that *C. d. rhodostictus* was highly paraphyletic. In addition to clusters 6 and 7, Admixture found support for another cluster 8) in the Mojave Desert. The tree shows a successive pattern of divergence from the Mojave into the Great Basin of Nevada, and with the most deeply-nested individuals occurring at the temperate-zone northern and northeastern range limits, including the northernmost described subspecies, *C. d. myurus* (Richardson, 1915). Admixture also identified 9) *C. d. myurus* as a cluster, albeit one that intergrades broadly with *C. d. rhodostictus* in the Mojave Desert. Another lineage includes individuals progressively nested east, then south, through the Salton Trough into the core of the Sonoran Desert. A long branch leads to a strongly supported clade containing lizards from four other previously named subspecies not yet discussed in this paper: *C. d. ventralis* (Hallowell, 1852) from the Yuman and Arizona upland subdivisions of the Sonoran Desert, following the Gila River to the range limit in southwestern New Mexico; *C. d. inusitatus* (Dickerson, 1919) from central Sonora and the Sonoran Gulf coast as far south as Guaymas, including Isla Tiburón; *C. d. brevipes* (Bogert & Dorson, 1942) from the subtropical dry forests of southern Sonora; and *C. d. bogerti* (Martin del Campo, 1943) from the tropical Sinaloan coast. The Admixture analysis identified 9) *C. d. ventralis*, 10) *C. d. inusitatus*, and 11) *C. d. bogerti* as distinct clusters under K=12, while the two individuals of *C. d. brevipes* are depicted as admixed between *C. d. bogerti* (larger contribution) and *C. d. inusitatus*.

**Figure 9.**
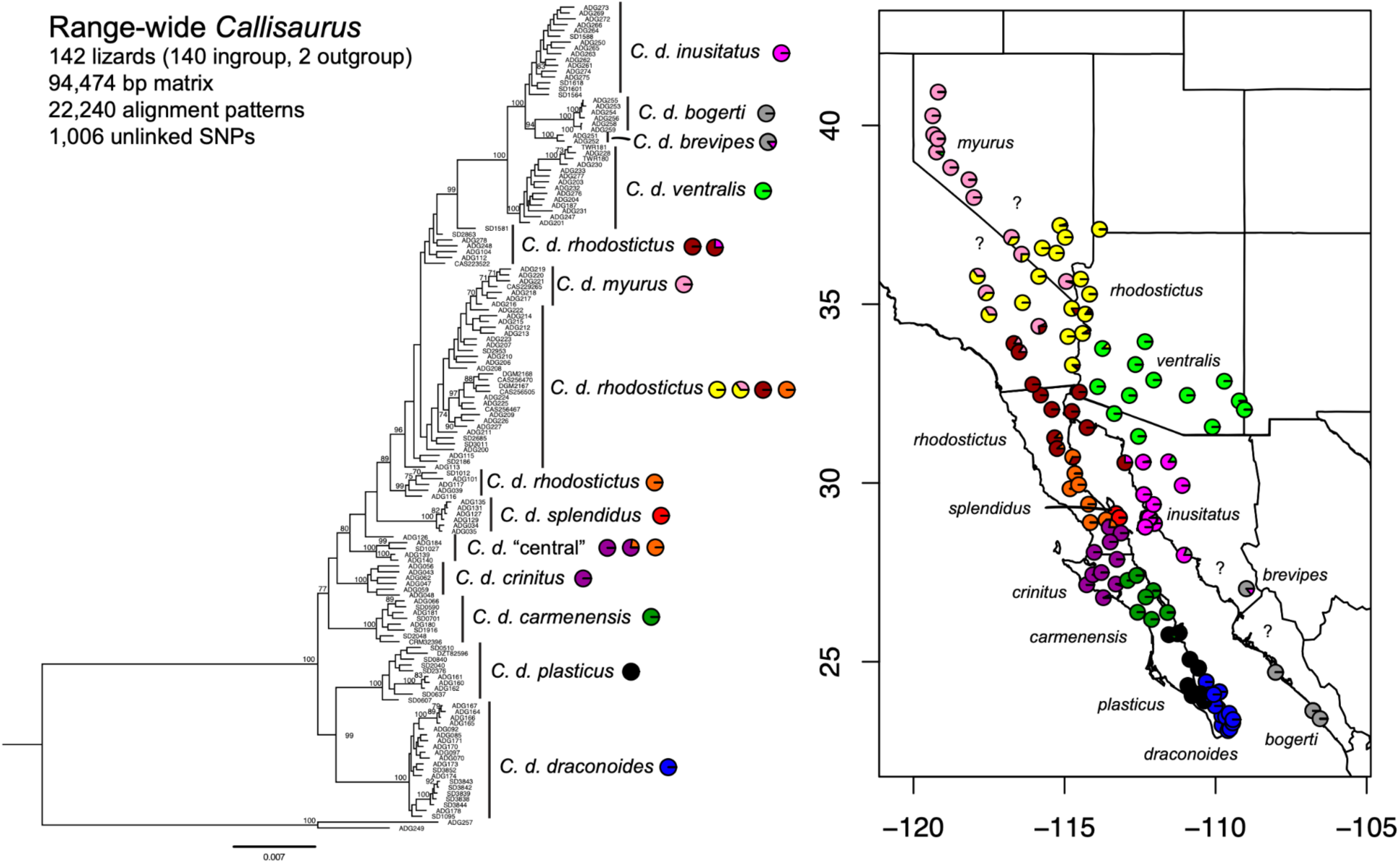
Maximum likelihood phylogram of concatenated ddRAD data (94,474 bp matrix, 22,240 alignment patterns) for *Callisaurus draconoides* across its range (n=140), rooted with *Holbrookia elegans* (n=2). Clades with bootstrap support > 70% are shown on nodes. The map on the right represents the Admixture results for 1,006 unlinked SNPs (optimal K=12), also annotated on the tree. Question marks (?) indicate areas where further field sampling is likely to yield insights on population boundaries and zones of admixture.

## DISCUSSION

We identified the geographic distributions of evolutionarily significant lineages across the BCP with relatively dense geographic sampling and shared phylogeographic and population genetic discontinuities across species. In the following sections we discuss the biogeographic patterns identified, including dispersal and vicariance, across tectonically active landscapes at the Isthmus of La Paz, Vizcaíno Desert, and the Gulf of California Islands. We then discuss the recent expansion of *Callisaurus* from the peninsula into the continental interior and taxonomic implications for all five species complexes.

### Isthmus of La Paz

The Isthmus of La Paz is one of two low-lying topographic breaks in the otherwise continuous peninsular ranges (the other being the Vizcaíno Desert). At its narrowest, the Isthmus is ∼42 km wide with a subtle crest at ∼300 m elevation, and it separates the Sierra de la Giganta and Magdalena Plain from the Cape District and Sierra de la Laguna, each with distinct endemic species, vegetation, and physiography (Morrone, 2021). Transpeninsular seaways across the Isthmus of La Paz of late Miocene, Pliocene, or Plio-Pleistocene age have been proposed based on earlier phylogeographic studies of *Callisaurus* (Lindell et al., 2005), *Urosaurus* (Aguirre et al., 1999; Lindell et al., 2008), and other terrestrial vertebrates (Blair et al., 2009; Riddle et al., 2000). Although an especially warm period of the Pliocene 2.7–3.2 Ma when global sea levels were ∼22 m higher than present (K. G. Miller et al., 2012) might have inundated much of the Isthmus, and Pliocene marine fossils are known from low-lying areas of the Cape District (W. Miller, 1980), direct geological evidence of a transpeninsular seaway here remains elusive (Dolby et al., 2015).

We confirmed lineage divergence across the Isthmus in four of five co-distributed species complexes (we did not find a break here in *S. zosteromus*). 1) Within *Callisaurus*, which occupies the alluvial terrain of the Isthmus, the contact zone is sharp between *C. d. draconoides* and *C. d. plasticus*, and closely follows the La Paz fault. This divergence was dated to the late Miocene to early Pleistocene (2.3–5.92 Ma), consistent with the Plio-Pleistocene estimate from (Lindell et al., 2005), and no admixture was detected. 2) *Petrosaurus,* whose members are strictly saxicolous, is absent from the Isthmus itself and is restricted to rugged terrain in the Cape District (*P. thalassinus*) and the Sierra de la Giganta (*P. repens*). The distinctiveness of these species was confirmed, and their divergence was dated to the Plio-Pleistocene (1.39– 3.98 Ma). 3) *Urosaurus* is widespread throughout the Isthmus. We found deep divergence, late Miocene to Pliocene (3.37–8.9 Ma), with no detected admixture between *U. nigricauda* of the Cape + Magdalena Plains districts and *U. microscutatus* from the Sierra de la Giganta and more northerly peninsular ranges. As with *Callisaurus*, our estimated divergence dates are consistent with published estimates from mtDNA of late Miocene divergence across the Isthmus in peninsular *Urosaurus* (Lindell et al., 2008). 4) We also found Miocene to early Pliocene (4.18– 10.7 Ma) divergence between *S. hunsakeri* and *S. orcutti*, which are absent from the Isthmus itself and the Magdalena Plain. Within each of these four complexes, the Isthmus of La Paz divergences predate those across the Vizcaíno Desert, consistent with the findings of Riddle et al. (2000). Our ecoevolity analyses found that at least three separate divergence events are required to explain our data, and the inferred order of divergences (Fig. 8) largely mirrors those in the species tree. To summarize, we interpret these results as rejecting the model of a single vicariant event causing divergence across all four species complexes simultaneously at the Isthmus of La Paz. In addition, evidence suggesting a shared event is inconsistent across taxa for the two data sets.

### Vizcaíno Desert

The second region where an ancient transpeninsular seaway has been proposed is the Vizcaíno Desert (Dolby et al., 2015), considered a unique district within the Baja California biogeographic province (Morrone, 2021). The prominent westernmost headland is Punta Eugenia, which is aligned with the Molokai Fracture Zone, extending thousands of kilometers into the Pacific plate and delimiting geomorphic provinces with distinct tectonic histories along the west coast of North America (Gottscho, 2016). We found strong evidence for clusters or clades that meet somewhere in the Vizcaíno Desert in all five species complexes. 1) *Callisaurus* is unique among the taxa studied in that we found three lineages meeting here: *C. d. carmenensis*, *C. d. crinitus*, and *C. d.* “central”. No admixture was found between these three lineages. The *C. d. carmenensis* and *C. d.* “central” lineages meet somewhere in the Sierra de San Francisco, while *C. d. crinitus* is unique to this study in that it is the only lineage endemic to the sandy western desert and has specific adaptations (elongated toe fringes) to improve locomotion on this substrate (Luke, 1986). The divergence of these three lineages dates to the Plio-Pleistocene (1.19–3.31 Ma), considerably younger than the late Miocene to early Pliocene age proposed by Lindell et al. (2005). 2) We inferred a Pleistocene (0.59–2.19 Ma) divergence between northern and central populations of *P. repens*, a novel finding of this study, although there is a large sampling gap; the break occurs somewhere in the Sierra de la Libertad. 3) We estimate that northern and southern clusters of *U. microscutatus* diverged in the Plio-Pleistocene (1.1–4.09 Ma), with some admixture in the Sierra de San Francisco. This date is considerably younger than the late Miocene date inferred from mtDNA (Lindell et al., 2008), and instead overlaps with the earlier Pleistocene date based on allozyme data (Aguirre et al., 1999). 4) Northern and southern clades of *S. orcutti* meet somewhere in the Sierra de la Libertad; the sampling gap is large between Cataviña and the Sierra de San Francisco. This divergence was dated to the Plio-Pleistocene (0.61–3.27 Ma). 5) Northern and southern clades of *S. zosteromus* (*S. z. rufidorsum* and *S. z. zosteromus*), which diverged in the Plio-Pleistocene (1.35–4.88 Ma), meet in the southern Vizcaíno Desert near Laguna San Ignacio. Because there were three pairwise comparisons for *Callisaurus*, a total of seven pairwise divergences were evaluated with ecoevolity. At least three divergence events are required to explain these data in the ecoevolity results, thus rejecting the simplest model of a single vicariance event across all taxa studied. However, in the species tree analysis, there is a window of 1.25–2.19 Ma that falls in the 95% confidence intervals for all five population pairs. We are not the first authors to find support for multiple divergence events in the Vizcaíno Desert, as analysis of mtDNA from twelve species of small mammals and reptiles found support for two divergence events here (Leaché et al., 2007).

### Gulf and Pacific Islands

For the amphibians and reptiles of the BCP, the Gulf Islands district contains the most species (84) and the most peninsular endemics (50) and therefore is of the greatest conservation significance (Peralta-García et al., 2023). In this study, we included 40 specimens (26 of which were *Callisaurus*) from nine islands in the Gulf and one in the Pacific (Isla Santa Margarita). These islands can be divided into three types based on their historical land-sea interactions. 1) Land-bridge islands are separated by shallow seaways that would have been exposed during Pleistocene glacial maxima when sea levels were up to 100 m lower than present. Islas Santa Margarita, Espiritu Santo, Partida, San Francisco, Danzante, Carmen, and Tiburón fall into this category (Carreño & Helenes, 2002). As expected, for these islands, the lizards we sampled always group with clusters and clades from the adjacent mainland, although some constitute monophyletic groups, e.g., *Callisaurus* from Islas Espiritu Santo and San Francisco, while others do not, e.g., *Callisaurus* from Isla Tiburón. 2) By contrast, oceanic islands are volcanoes that emerge from the seafloor and were never connected to a continent. It can thus be inferred that any lizards occupying these islands must have dispersed over water (or were introduced by humans). The only oceanic island in our study is Isla Tortuga, a caldera of Pleistocene-Holocene age emerging from waters >1000 m deep (Carreño & Helenes, 2002). Its population of *S. orcutti* (samples 39–40) is sister to but assigned to the same cluster as the southern peninsular *S. orcutti*. As with the land-bridge islands, we did not explicitly date the divergence of the Isla Tortuga population, but it is shallower in the tree than the Vizcaíno split mentioned above, which was dated to the Plio-Pleistocene. 3) Deep-water islands composed of continental rock, Islas Cerralvo and Ángel de la Guarda, do not neatly fit into the previous categories. Isla Cerralvo is isolated from the rest of the Cape Region by the Canal de Cerralvo (>200 m depth), and the mode of origin is thought to be uplift (Carreño & Helenes, 2002). Samples of *C. d. draconoides* from Isla Cerralvo were included in the range-wide *Callisaurus* tree (SD3838, 3839, 3842–3844), where they form a clade within *C. d. draconoides* with a branch length comparable to that of the Isla Espiritu Santo clade (ADG164–167). Isla Ángel de la Guarda, the second-largest island in the Gulf (931 km^2^) is also the only island in this study with relatively precise dating in both our dataset and the geological literature. Isla Ángel de la Guarda separated from the BCP during the Plio-Pleistocene (2–3.3 Ma) due to transform faulting and rifting, forming a deep basin (>1000 m depth) with an active plate boundary in the Canal de Ballenas (Aragón-Arreola & Martín-Barajas, 2007; Dolby et al., 2015; Nagy & Stock, 2000). This estimate overlaps partially with the 95% HPD of 0.66–2.36 Ma for the divergence of *C. d. splendidus* (the only island population in the BCP ddRAD dataset to be recognized as a distinct cluster by Admixture) and *C. d. rhodostictus*. Thus, both early Pleistocene (2–2.36 Ma) vicariance and late Pleistocene (0.66–2 Ma) dispersal origins of *C. d. splendidus* remain plausible. *Petrosaurus slevini*, as traditionally circumscribed, is also endemic to Isla Ángel de la Guarda, but our analyses grouped this population with southern *P. mearnsi* (together designated here as *P. “slevini”*), although we did not explicitly date the divergence of the island from peninsular populations, we did date the *P. mearnsi / P. “slevini”* split at 2.49 Ma (95% HPD 1.02–4.22 Ma), and the colonization of the island must be younger than this. To summarize, our results are generally consistent with published geological ages for all of the islands studied, with *C. d. splendidus* on Isla Ángel de la Guarda being the most divergent.

### Historical biogeography of *Callisaurus*

We collected a range-wide ddRAD dataset to test three hypotheses about the historical biogeography of *Callisaurus*. 1) The vicariance hypothesis predicts a sister relationship between divergent mainland and peninsular lineages. Examples of lizards demonstrating this pattern in the BCP vs. continent include *S. zosteromus* and *S. magister* (this study; Grismer & McGuire, 1996; Leaché & Mulcahy, 2007; Pavón-Vázquez et al., 2024; Schulte et al., 2006), *Crotaphytus* (McGuire et al., 2007), *Coleonyx* (Leavitt et al., 2020) and *Sauromalus* (Sumarli et al., 2024). 2) The Gulf of California is ∼200 km wide at its broadest, but in the “Midriff” region it is much narrower, and a string of islands, spaced no more than 17 km apart, make it possible for some taxa to cross the Gulf in either direction via stepping-stone dispersal. This pattern is most commonly seen in plants, especially those that rely on bats for pollination (Arenas et al., 2023; Cody et al., 2002), but also in other groups, e.g., *Phyllodactylus* geckos (Murphy & Aguirre-León, 2002). To test this hypothesis for *Callisaurus* requires samples from Islas Ángel de la Guarda and Isla Tiburón, the two largest Midriff islands, which are only 53 km apart, and the adjacent peninsula and continent. The stepping-stone hypothesis predicts that lizards from three of the populations, either the peninsular or the mainland population and the two island populations (in order of their geographic proximity), are successively more closely related to the fourth. 3) The third hypothesis is overland dispersal and range expansion from the mainland to the peninsula, or vice versa. Under this scenario, lizards evolved on either the mainland or the peninsula, and then traveled the long way around the Gulf of California instead of dispersing over water barriers or hitching a ride on the Pacific Plate. This hypothesis predicts that populations representing recent range expansions are nested within those representing the area of origin.

The *Callisaurus* phylogeny favors the third hypothesis, rejecting the Gulf vicariance and Midriff stepping-stone models. The divergence between *C. d. splendidus* and *C. d. rhodostictus* from northern Baja California dates to the Pleistocene (95% HPD = 0.66–2.36 Ma), constraining when range expansion could have occurred. The expansion out of northern Baja (Figure 9) progresses in a south-to-north pattern of sequential cladogenesis after this time. We hypothesize that as the northern Gulf receded from the San Gorgonio Pass and Salton Trough at the end of the Pliocene, this opened a dispersal corridor into the mainland. From there, *Callisaurus* expanded its range across western North America into temperate, subtropical, and tropical biomes. To the best of our knowledge, this peninsula-to-mainland pattern of expansion has not been documented before. Reptiles and plant taxa that have been shown to have south-to-north range expansions within the BCP, or expansions into the California floristic province, include *Crotaphytus vestigium* (McGuire et al., 2007), *Euphorbia lomelii* (Garrick et al., 2009), *Crotalus ruber* (Harrington et al., 2018), *Dipsosaurus dorsalis* (Valdivia-Carrillo et al., 2017), *Phrynosoma coronatum* species complex (Leaché et al., 2009), and *U. microscutatus* (this study), but these examples are all within the BCP. *Hypsiglena* has been shown to demonstrate both southern vicariance and northern dispersal, forming a ring distribution around the Gulf (Mulcahy & Macey, 2009), but the direction of northward dispersal was from mainland to peninsula. Therefore, the multi-directionality, scale and recency of the continental expansion of *Callisaurus*—the range front, from the Black Rock Desert to the Sinaloan coast, ∼2,300 km wide—is perhaps the most novel finding of this study. As Riddle et al. (2000) concluded, most of the “phylogeographic architecture” of the peninsula is “cryptically embedded within widespread taxonomic species and species-groups, such that the unique evolutionary history of the Peninsular Desert has been obscured and ignored.” This statement certainly applies to *Callisaurus,* although the subspecies revealed it to some degree. In summary, the recent expansion, and its classification as a monospecific genus, has until now masked that *Callisaurus* likely evolved as a peninsular endemic, and still contains several lineages endemic to the BCP.

### Taxonomic implications

Adest (1987) used allozymes to study geographic variation within *C. draconoides*, found little genetic differentiation over the range, and concluded that *Callisaurus* is best treated as monotypic. Grismer (2002) also considered *C. draconoides* in the BCP a single species, but divided into five “pattern classes” (which we treat here as subspecies, as they were previously treated) within the BCP. The seven peninsular lineages revealed by the BCP ddRAD analyses are a close fit to the current subspecies, with the following exceptions. The *C. d.* “central” clade corresponds to an intergrade zone in Grismer’s map between *C. d. rhodostictus* x *crinitus* x *carmenensis*, and *C. d. plasticus* from our analyses roughly matches Grismer’s intergrade between *C. d. draconoides* x *carmenensis,* except that the range of *C. d. plasticus* includes the southern part of the distribution of Grismer’s *carmenensis*. When mainland samples are included, the distributions of monophyletic lineages of *Callisaurus* include specimens from, or very close to, type localities for *C. d. ventralis*, *C. d. brevipes*, and *C. d. bogerti*. *C. d. rhodostictus* is highly paraphyletic with respect to remaining mainland lizards. *Callisaurus d. myurus* is monophyletic but deeply nested within *C. d. rhodostictus* and exhibits considerable admixture with *C. d. rhodostictus* in the Mojave Desert. Therefore, we provisionally recognize the eleven previously named subspecies of *C. draconoides* shown in Figure 9, although *C. d. rhodostictus* is likely composed of multiple lineages, and *C. d. bogerti* and *C. d. brevipes* may form a single lineage, so that all three taxa should be considered questionable (de Queiroz & Chan, 2025).

Within *Petrosaurus*, *P. slevini* has been recognized as a distinct species from *P. mearnsi* based on color, throat pattern, and body size (Grismer, 1999, 2002), but our phylogenetic analyses found that *P. slevini* is nested within *P. mearnsi,* which was instead divided into northern and southern lineages that do not correspond to currently recognized taxa. The northern lineage, which includes the type locality for *P. mearnsi*, ranges from San Gorgonio Pass to the Sierra San Pedro Martír, while the southern lineage includes *P. mearnsi* from Cataviña to Bahía de los Ángeles and *P. slevini* from Isla Ángel de la Guarda. Based on limited sampling between Bahía de los Ángeles and the Sierra San Pedro Martír, we do not propose changes to *Petrosaurus* taxonomy at this time.

Within *Urosaurus*, Aguirre et al. (1999) unified *U. microscutatus* with *U. nigricauda* because patterns of allelic distribution suggested ongoing gene flow. Grismer (2002) followed this taxonomy. Although we found support for some introgression at the contact zone in a single individual, in our study these lineages were found to be reciprocally monophyletic and relatively ancient, diverging in the late Miocene/Pliocene. We therefore resurrect the species *U. microscutatus*. We do not recognize formal taxa for northern and southern populations within *U. microscutatus* because of the sampling gap between the Sierra de la Liberdad and the Sierra de San Francisco and evidence of admixture in several of the samples.

Although both *S. orcutti* and *S. zosteromus* have northern and southern lineages, significant sampling gaps remain in the northern BCP for *S. orcutti*. The northern lineage of *S. zosteromus* corresponds to the previously recognized “*rufidorsum*” plus a putative intergrade zone (Grismer, 2002), while the southern lineage corresponds to the combined “*zosteromus*” and “*monserratensis*” (Grismer, 2002). Our results thus corroborate the division of *S. zosteromus* into northern and southern lineages (Pavón-Vázquez et al., 2024) and narrow the sampling gap between them. We therefore recognize the subspecies *S. z. rufidorsum* for the northern lineage and *S. z. zosteromus* for the southern one, although further study is needed to assess the degree of separation between those lineages. The current lack of data for specimens from Isla Monserrat (the type locality of *S. z. monserratensis*) leaves the taxonomic status of that population uncertain.

For all taxa, future research should explicitly test alternative species delimitation hypotheses in a model-based framework, and all data from this study are being made publicly available to that end.

### Conclusions

Among five co-distributed complexes of phrynosomatid lizards we examined within the BCP, four exhibited contact zones between lineages at the Isthmus of La Paz, and all five did in the Vizcaíno Desert. Our results require at least three separate divergence events across both biogeographic barriers, and for each, we reject the simplest vicariance hypothesis of a single episode of divergence across all taxa. In the Gulf of California, our divergence time estimate for *Callisaurus* on Isla Ángel de la Guarda partially overlaps with when the island separated from the peninsula, consistent with both vicariance and dispersal scenarios, adding confidence to our timetree. A clade of *S. orcutti* on the oceanic Isla Tortuga requires an overwater dispersal origin. South-to-north range expansions within the peninsula are evident in *Urosaurus* and *Callisaurus.* A novel finding of our study is a Pleistocene range expansion of *Callisaurus* from the peninsula into the continental mainland that allowed *Callisaurus* to colonize novel temperate, subtropical, and tropical biomes, with a range front of ∼2,300 km. This expansion has obscured that *Callisaurus* likely was a peninsular endemic in the late Miocene and Pliocene and still contains several lineages endemic to the BCP. The current taxonomy also obscures these lineages, and based on our results, we propose minor taxonomic changes to better reflect evolutionary history. These findings collectively highlight the importance of the BCP’s tectonic isolation as a driver not only of local or peninsular endemism, but potentially also as a contributing factor to lineage diversification more broadly in the region.

## DATA ACCESSIBILITY

Raw sequence data have been deposited in the NCBI SRA (https://www.ncbi.nlm.nih.gov/sra/PRJNA1242740). Additional data and code necessary to reproduce the results presented here will be made available as a Dryad dataset upon acceptance of this manuscript to a peer-reviewed journal.

## AUTHOR CONTRIBUTIONS

A.D.G., B.D.H., J.L.E., T.W.R., and K.D.Q. designed the study; A.D.G., B.D.H., and J.L.E. planned and executed the field collection of specimens and their subsequent deposition into museums; A.D.G. generated molecular data in the laboratories of T.W.R. and K.D.Q.; A.D.G. and A.D.L analyzed the data and made the figures. All authors contributed to interpreting the results and writing the manuscript.

## ACKNOWLEDGMENTS

For funding, we thank the University of California Institute for Mexico and the United States (Dissertation Research Grant), the Anza-Borrego Foundation (Howie Wier Memorial Conservation Grant), the Community Foundation (Desert Legacy Fund), the American Philosophical Society (Lewis and Clark Fund For Exploration and Field Research), the National Science Foundation (Doctoral Dissertation Improvement Grant, Award Number 1406589 to A.D.G. and T.W.R., NSF-SBS-2023723 to A.D.L.), and the Smithsonian Institution (Peter Buck Postdoctoral Fellowship to A.D.G.). Permits to collect animals were provided by the California Department of Fish and Wildlife (SC-9768), Arizona Game and Fish Department (SP698679, SP733215), Nevada Department of Wildlife (491604), and SEMARNAT (SGPA/DGVS/01239/12, SGPA/DGVS/00916/13, SGPA/DGVS/00976/14, SGPA/DGVS/01207/16). Export permits from Mexico to the U.S. were provided by PROFEPA (2012-30461, 2013-32393, 2014-34440, 2016-39585), and importations were approved by the U.S. Fish and Wildlife Service. Animal welfare protocols were approved by San Diego State University (APF# 12-04-010R, 14-03-008R) and the Smithsonian Institution (ACUC# 2016-01). The 2003 binational expedition to the Agua Verde-Punta Mechudo conservation corridor, including O. Flores-Villela, J. H. Valdez Villavicencio, A. Peralta-Garcia, D. A. Wood, C. Jauregui, T. Myers, and R. Hill, contributed to the collection of specimens used in this study. We are grateful to other individuals who assisted with collecting and preparing specimens: G. Ruiz-Campos, J. Rabbers, S. Harrington, N. Angeli, M. Ziebell, F. E. B. Herrera, R. Bell, A. Stubbs, D. Mulcahy, J. Andermann, R. A. Gottscho, A. Meier, S. Murray, H. Heinz, P. Maier, T. Higham, C. Collins, D. Leavitt, M. Reinbold, N. Marshall, T. Devitt, J. McGuire, C. Mahrdt, B. Lowe, T. Townsend, and A. Marion. We thank the Comcáac people for granting us permission to collect on tribal territory (Isla Tiburón). For assistance with laboratory work we especially thank J. Grummer, P. Maier, L. Dinsdale, F. Rohwer, M. Braun and N. White. Most analyses in this paper were conducted on the Smithsonian High Performance Cluster (SI/HPC), Smithsonian Institution (https://doi.org/10.25572/SIHPC), and we thank M. Lloyd, V. Gonzalez, R. Dikow, and M. Kweskin for thoughtful advice and troubleshooting.

## SUPPLEMENTAL

**Supplemental Figure 1.** Concatenated TSC tree (BEAST)

**Supplemental Figure 2.** Range-wide *Callisaurus* tree with Admixture results

**Supplemental Table 1.** Specimen information, BCP ddRAD

**Supplemental Table 2.** Specimen information, *Callisaurus* ddRAD

**Supplemental Table 3.** Specimen information, TSC

**Supplemental Table 4.** Summary statistics, BCP ddRAD

**Supplemental Table 5.** Summary statistics, TSC

## REFERENCES

Adest, G. A. (1987). Genetic Differentiation among Populations of the Zebratail Lizard, *Callisaurus draconoides* (Sauria: Iguanidae). Copeia, 1987(4), 854–859. 10.2307/1445547

Aguirre, G., Morafka, D., & Murphy, R. (1999). The peninsular archipelago of Baja California: A thousand kilometers of tree lizard genetics. Herpetologica, 55(3), 369–381.

Alexander, D. H., Novembre, J., & Lange, K. (2009). Fast model-based estimation of ancestry in unrelated individuals. Genome Research, 19(9), 1655–1664. 10.1101/gr.094052.109

Alföldi, J., Di Palma, F., Grabherr, M., Williams, C., Kong, L., Mauceli, E., Russell, P., Lowe, C. B., Glor, R. E., Jaffe, J. D., Ray, D. A., Boissinot, S., Shedlock, A. M., Botka, C., Castoe, T. A., Colbourne, J. K., Fujita, M. K., Moreno, R. G., ten Hallers, B. F.,… Lindblad-Toh, K. (2011). The genome of the green anole lizard and a comparative analysis with birds and mammals. Nature, 477(7366), 587–591. 10.1038/nature10390

Aragón-Arreola, M., & Martín-Barajas, A. (2007). Westward migration of extension in the northern Gulf of California, Mexico. Geology, 35(6), 571–574. 10.1130/G23360A.1

Arenas, S., Búrquez, A., Bustamante, E., Scheinvar, E., & Eguiarte, L. E. (2023). Are 150 km of open sea enough? Gene flow and population differentiation in a bat-pollinated columnar cactus. PLOS ONE, 18(6), e0282932. 10.1371/journal.pone.0282932

Bennett, S. E. K., & Oskin, M. E. (2014). Oblique rifting ruptures continents: Example from the Gulf of California shear zone. Geology, 42(3), 215–218. 10.1130/G34904.1

Blainville, H. (1835). Description de quelques espèces de reptiles de la Californie précédée de l’analyse d’un système général d’herpétologie et d’amphibologie. Nouvelles Annales Du Muséum d’histoire Naturelle Paris, 4, 232–296.

Blair, C., Méndez de la Cruz, F. R., Ngo, A., Lindell, J., Lathrop, A., & Murphy, R. W. (2009). Molecular phylogenetics and taxonomy of leaf-toed geckos (Phyllodactylidae: *Phyllodactylus*) inhabiting the peninsula of Baja California. Zootaxa, 2027(1), 28–42. 10.11646/zootaxa.2027.1.2

Bogert, C., & Dorson, E. (1942). A new lizard of the genus *Callisaurus* from Sonora. Copeia, 1942(3), 173–175.

Bouckaert, R., Heled, J., Kühnert, D., Vaughan, T., Wu, C.-H., Xie, D., Suchard, M. A., Rambaut, A., & Drummond, A. J. (2014). BEAST 2: A Software Platform for Bayesian Evolutionary Analysis. PLOS Computational Biology, 10(4), e1003537. 10.1371/journal.pcbi.1003537

Carreño, A. L. (1985). Biostratigraphy of the late Miocene to Pliocene on the Pacific island Maria Madre, Mexico. Micropaleontology, 31(2), 139–166.

Carreño, A. L. (1992). Neogene microfossils from the Santiago Diatomite, Baja California Sur, Mexico. Revista Paleontología Mexicana, 59, Article 59. 10.22201/igl.05437652e.1992.0.59.274

Carreño, A. L., & Helenes, J. (2002). Geology and Ages of the Islands. In T. J. Case, M. L. Cody, & E. Ezcurra (Eds.), A new island biogeography of the Sea of Cortés (pp. 14–40). Oxford University Press.

Chang, C. C., Chow, C. C., Tellier, L. C., Vattikuti, S., Purcell, S. M., & Lee, J. J. (2015). Second-generation PLINK: Rising to the challenge of larger and richer datasets. GigaScience, 4(1), s13742–015-0047–0048. 10.1186/s13742-015-0047-8

Cody, M., Moran, R., Rebman, J., Thompson, H., & Robles, A. (2002). Plants. In T. J. Case, M. L. Cody, & E. Ezcurra (Eds.), A new island biogeography of the Sea of Cortés (pp. 63– 111). Oxford University Press.

Cope, E. D. (1863). Descriptions of New American Squamata, in the Museum of the Smithsonian Institution, Washington. Proceedings of the Academy of Natural Sciences of Philadelphia, 15, 100–106.

Cope, E. D. (1864). Contributions to the herpetology of tropical America. Proc. Acad. Nat. Sci. Philadelphia, 16, 166–181.

Cope, E. D. (1896). On the genus *Callisaurus*. American Naturalist, 30, 1049–1050.

Crews, S. C., & Hedin, M. (2006). Studies of morphological and molecular phylogenetic divergence in spiders (Araneae: *Homalonychus*) from the American southwest, including divergence along the Baja California Peninsula. Molecular Phylogenetics and Evolution, 38(2), 470–487. 10.1016/j.ympev.2005.11.010

de Queiroz, K., & Chan, L. M. (2025). Squamata (excluding snakes) – Lizards. In K. E. Nicholson (Ed.), Scientific and Standard English Names of Amphibians and Reptiles of North America North of Mexico, with Comments Regarding Confidence in Our Understanding (pp. 1–87).

Dickerson, M. C. (1919). Diagnoses of twenty-three new species and a new genus of lizards from Lower California. Bulletin of the AMNH, 61, 461–477.

Dolby, G. A., Adrian Munguia-Vega, Dorsey, R. J., Bennett, S. E. K., Darin, M., Gardner, K., Araya-Donoso, R., Davalos-Dehullu, L., Baty, S., Biddy, A., Andreev, V., Lira-Noriega, A., Wilder, B. T., Cortez, D., Culver, M., & Kusumi, K. (2024, September 22). A new working model for co-evolution of plant and animal species on the Baja California peninsula from genomic and geologic data. GSA Connects 2024 Meeting in Anaheim, California. https://gsa.confex.com/gsa/2024AM/meetingapp.cgi/Paper/404092

Dolby, G. A., Bennett, S. E. K., Lira-Noriega, A., Wilder, B. T., & Munguía-Vega, A. (2015). Assessing the Geological and Climatic Forcing of Biodiversity and Evolution Surrounding the Gulf of California. Journal of the Southwest, 57(2), 391–455.

Eaton, D. A. R. (2014). PyRAD: Assembly of de novo RADseq loci for phylogenetic analyses. Bioinformatics, 30(13), 1844–1849. 10.1093/bioinformatics/btu121

Faircloth, B. C. (2016). PHYLUCE is a software package for the analysis of conserved genomic loci. Bioinformatics, 32(5), 786–788. 10.1093/bioinformatics/btv646

Faircloth, B. C., McCormack, J. E., Crawford, N. G., Harvey, M. G., Brumfield, R. T., & Glenn, T. C. (2012). Ultraconserved elements anchor thousands of genetic markers spanning multiple evolutionary timescales. Systematic Biology, 61(5), 717–726. 10.1093/sysbio/sys004

Gardner, K., Hasiotis, S., Dorsey, R. J., Darin, M., Hausback, B., Bennett, S. E. K., Heizler, M., & Dolby, G. (2024). Evidence for Terrestrial Origin of the Pliocene San Regis Beds, Central Baja California Peninsula, Mexico. Geological Society of America Abstracts, 56, 401950. 10.1130/abs/2024AM-401950

Garrick, R. C., Nason, J. D., Meadows, C. A., & Dyer, R. J. (2009). Not just vicariance: Phylogeography of a Sonoran Desert euphorb indicates a major role of range expansion along the Baja peninsula. Molecular Ecology, 18(9), 1916–1931. 10.1111/j.1365-294X.2009.04148.x

Gastil, R. G. (1978). A reconnaissance geologic map of the west-central part of the state of Nayarit, Mexico [Map]. Geological Society of America, [1978] ©1978. https://search.library.wisc.edu/catalog/999506573602121

Gottscho, A. D. (2015). Lineage Diversification of Lizards (Phrynosomatidae) in Southwestern North America: Integrating Genomics and Geology [UC Riverside]. https://escholarship.org/uc/item/8nk5d7b1

Gottscho, A. D. (2016). Zoogeography of the San Andreas Fault system: Great Pacific Fracture Zones correspond with spatially concordant phylogeographic boundaries in western North America. Biological Reviews, 91(1), 235–254. 10.1111/brv.12167

Gottscho, A. D., Mulcahy, D. G., Leaché, A. D., de Queiroz, K., & Lovich, R. E. (2024). Population genomics of flat-tailed horned lizards (*Phrynosoma mcallii*) informs conservation and management across a fragmented Colorado Desert landscape. Molecular Ecology, 33(7), e17308. 10.1111/mec.17308

Gottscho, A. D., Wood, D. A., Vandergast, A. G., Lemos-Espinal, J., Gatesy, J., & Reeder, T. W. (2017). Lineage diversification of fringe-toed lizards (Phrynosomatidae: *Uma notata* complex) in the Colorado Desert: Delimiting species in the presence of gene flow. Molecular Phylogenetics and Evolution, 106, 103–117. 10.1016/j.ympev.2016.09.008

Grismer, L. (1999). An Evolutionary Classification of Reptiles on Islands in the Gulf of California, México. Herpetologica, 55(4), 446–469.

Grismer, L. (2002). Amphibians and Reptiles of Baja California Including its Pacific Islands and the Islands in the Sea of Cortés. University of California Press.

Grismer, L., & McGuire, J. A. (1996). Taxonomy and Biogeography of the *Sceloporus magister* Complex (Squamata: Phrynosomatidae) in Baja California, México. Herpetologica, 52(3), 416–427.

Hall, W., & Smith, H. (1979). Lizards of the *Sceloporus orcutti* complex of the Cape Region of Baja California. Breviora, 452, 1–26.

Hallowell, E. (1852). Descriptions of new species of reptiles inhabiting North America. Proc. Acad. Nat. Sci. Philadelphia, 6, 177–182.

Hallowell, E. (1854). Descriptions of new reptiles from California. Proceedings of the Academy of Natural Sciences of Philadelphia, 7, 91–97.

Harrington, S. M., Hollingsworth, B. D., Higham, T. E., & Reeder, T. W. (2018). Pleistocene climatic fluctuations drive isolation and secondary contact in the red diamond rattlesnake (*Crotalus ruber*) in Baja California. Journal of Biogeography, 45(1), 64–75. 10.1111/jbi.13114

Leaché, A. D., Chavez, A. S., Jones, L. N., Grummer, J. A., Gottscho, A. D., & Linkem, C. W. (2015). Phylogenomics of phrynosomatid lizards: Conflicting signals from sequence capture versus restriction site associated DNA sequencing. Genome Biology and Evolution, 7(3), 706–719. 10.1093/gbe/evv026

Leaché, A. D., Crews, S. C., & Hickerson, M. J. (2007). Two waves of diversification in mammals and reptiles of Baja California revealed by hierarchical Bayesian analysis. Biology Letters, 3(6), 646–650. 10.1098/rsbl.2007.0368

Leaché, A. D., Koo, M. S., Spencer, C. L., Papenfuss, T. J., Fisher, R. N., & McGuire, J. A. (2009). Quantifying ecological, morphological, and genetic variation to delimit species in the coast horned lizard species complex (*Phrynosoma*). Proceedings of the National Academy of Sciences, 106(30), 12418–12423. 10.1073/pnas.0906380106

Leaché, A. D., & Mulcahy, D. G. (2007). Phylogeny, divergence times and species limits of spiny lizards (*Sceloporus magister* species group) in western North American deserts and Baja California. Molecular Ecology, 16(24), 5216–5233. 10.1111/j.1365-294X.2007.03556.x

Leavitt, D. H., Hollingsworth, B. D., Fisher, R. N., & Reeder, T. W. (2020). Introgression obscures lineage boundaries and phylogeographic history in the western banded gecko, *Coleonyx variegatus* (Squamata: Eublepharidae). Zoological Journal of the Linnean Society, 190(1), 181–226. 10.1093/zoolinnean/zlz143

Lin, H.-C., Sánchez-Ortiz, C., & Hastings, P. A. (2009). Colour variation is incongruent with mitochondrial lineages: Cryptic speciation and subsequent diversification in a Gulf of California reef fish (Teleostei: Blennioidei). Molecular Ecology, 18(11), 2476–2488. 10.1111/j.1365-294X.2009.04188.x

Lindell, J., Méndez-de la Cruz, F. R., & Murphy, R. W. (2005). Deep genealogical history without population differentiation: Discordance between mtDNA and allozyme divergence in the zebra-tailed lizard (*Callisaurus draconoides*). Molecular Phylogenetics and Evolution, 36(3), 682–694. 10.1016/j.ympev.2005.04.031

Lindell, J., Mendez-De La Cruz, F. R., & Murphy, R. W. (2008). Deep biogeographical history and cytonuclear discordance in the black-tailed brush lizard (*Urosaurus nigricaudus*) of Baja California. Biological Journal of the Linnean Society, 94(1), 89–104. 10.1111/j.1095-8312.2008.00976.x

Lindell, J., Ngo, A., & Murphy, R. W. (2006). Deep genealogies and the mid-peninsular seaway of Baja California. Journal of Biogeography, 33(8), 1327–1331. 10.1111/j.1365-2699.2006.01532.x

Lischer, H. E. L., & Excoffier, L. (2012). PGDSpider: An automated data conversion tool for connecting population genetics and genomics programs. Bioinformatics, 28(2), 298–299. 10.1093/bioinformatics/btr642

Lonsdale, P. (1989). Geology and tectonic history of the Gulf of California. In E. L. Winterer, D. M. Hussong, & R. W. Decker (Eds.), The Eastern Pacific Ocean and Hawaii: Vol. N (p. 0). Geological Society of America. 10.1130/DNAG-GNA-N.499

Luke, C. (1986). Convergent evolution of lizard toe fringes. Biological Journal of the Linnean Society, 27(1), 1–16. 10.1111/j.1095-8312.1986.tb01723.x

Martin del Campo, R. (1943). *Callisaurus draconoides bogerti* subsp. Nov. Ann. Inst. Biol. Mexico, 14, 619–621.

McCloy, C., Ingle, J. C., & Barron, J. A. (1988). Neogene stratigraphy, foraminifera, diatoms, and depositional history of Maria Madre Island, Mexico: Evidence of early Neogene marine conditions in the southern Gulf of California. Marine Micropaleontology, 13(3), 193–212. 10.1016/0377-8398(88)90003-5

McGuire, J. A., Linkem, C. W., Koo, M. S., Hutchison, D. W., Lappin, A. K., Orange, D. I., Lemos-Espinal, J., Riddle, B. R., & Jaeger, J. R. (2007). Mitochondrial introgression and incomplete lineage sorting through space and time: Phylogenetics of crotaphytid lizards. Evolution, 61(12), 2879–2897. 10.1111/j.1558-5646.2007.00239.x

Miller, K. G., Wright, J. D., Browning, J. V., Kulpecz, A., Kominz, M., Naish, T. R., Cramer, B. S., Rosenthal, Y., Peltier, W. R., & Sosdian, S. (2012). High tide of the warm Pliocene: Implications of global sea level for Antarctic deglaciation. Geology, 40(5), 407–410. 10.1130/G32869.1

Miller, W. (1980). The late Pliocene Las Tunas local fauna from southernmost Baja California, Mexico. Journal of Paleontology, 54, 762–805.

Mittermeier, R. A., Mittermeier, C. G., Brooks, T. M., Pilgrim, J. D., Konstant, W. R., da Fonseca, G. a. B., & Kormos, C. (2003). Wilderness and biodiversity conservation. Proceedings of the National Academy of Sciences of the United States of America, 100(18), 10309–10313. 10.1073/pnas.1732458100

Molina-Cruz, A. (1994). Biostratigraphy and paleoceanographic significance of the radiolarians from the protomouth of the Gulf of California. Ciencias Marinas, 20(4), Article 4. 10.7773/cm.v20i4.980

Moore, D. G., & Buffington, E. C. (1968). Transform faulting and growth of the Gulf of California since the late Pliocene. Science (New York, N.Y.), 161(3847), 1238–1241. 10.1126/science.161.3847.1238

Morrone, J. J. (2021). Biogeographic regionalisation of the Baja California biogeographic province, Mexico: A review. Journal of Natural History, 55(5–6), 365–379. 10.1080/00222933.2021.1903111

Mulcahy, D. G., & Macey, J. R. (2009). Vicariance and dispersal form a ring distribution in nightsnakes around the Gulf of California. Molecular Phylogenetics and Evolution, 53(2), 537–546. 10.1016/j.ympev.2009.05.037

Murphy, R. (1983). Paleobiogeography and genetic differentiation of the Baja California herpetofauna. ResearchGate, 137, 1–48.

Murphy, R., & Aguirre-León, G. (2002). The Nonavian Reptiles: Origins and Evolution. In T. J. Case, M. L. Cody, & E. Ezcurra (Eds.), A new island biogeography of the Sea of Cortés (pp. 181–220). Oxford University Press.

Nagy, E. A., & Stock, J. M. (2000). Structural controls on the continent-ocean transition in the northern Gulf of California. Journal of Geophysical Research: Solid Earth, 105(B7), 16251–16269. 10.1029/1999JB900402

Oaks, J. R. (2019). Full Bayesian Comparative Phylogeography from Genomic Data. Systematic Biology, 68(3), 371–395. 10.1093/sysbio/syy063

Ogilvie, H. A., Bouckaert, R. R., & Drummond, A. J. (2017). StarBEAST2 Brings Faster Species Tree Inference and Accurate Estimates of Substitution Rates. Molecular Biology and Evolution, 34(8), 2101–2114. 10.1093/molbev/msx126

Oskin, M., & Stock, J. (2003a). Marine incursion synchronous with plate-boundary localization in the Gulf of California. Geology, 31(1), 23–26. 10.1130/0091-7613(2003)031<0023:MISWPB>2.0.CO;2

Oskin, M., & Stock, J. (2003b). Pacific–North America plate motion and opening of the Upper Delfín basin, northern Gulf of California, Mexico. GSA Bulletin, 115(10), 1173–1190. 10.1130/B25154.1

Pavón-Vázquez, C. J., Rana, Q., Farleigh, K., Crispo, E., Zeng, M., Liliah, J., Mulcahy, D., Ascanio, A., Jezkova, T., Leaché, A. D., Flouri, T., Yang, Z., & Blair, C. (2024). Gene Flow and Isolation in the Arid Nearctic Revealed by Genomic Analyses of Desert Spiny Lizards. Systematic Biology, 73(2), 323–342. 10.1093/sysbio/syae001

Peralta-García, A., Valdez-Villavicencio, J. H., Fucsko, L. A., Hollingsworth, B. D., Johnson, J. D., Mata-Silva, V., Rocha, A., DeSantis, D. L., Porras, L. W., & Wilson, L. D. (2023). The herpetofauna of the Baja California Peninsula and its adjacent islands, Mexico: Composition, distribution, and conservation status. Amphibian & Reptile Conservation, 17(1), 57–142. 10.5281/zenodo.12762042

Peterson, B. K., Weber, J. N., Kay, E. H., Fisher, H. S., & Hoekstra, H. E. (2012). Double Digest RADseq: An Inexpensive Method for De Novo SNP Discovery and Genotyping in Model and Non-Model Species. PLOS ONE, 7(5), e37135. 10.1371/journal.pone.0037135

Peterson, D. L., Kubow, K. B., Connolly, M. J., Kaplan, L. R., Wetkowski, M. M., Leong, W., Phillips, B. C., & Edmands, S. (2013). Reproductive and phylogenetic divergence of tidepool copepod populations across a narrow geographical boundary in Baja California. Journal of Biogeography, 40(9), 1664–1675. 10.1111/jbi.12107

Rau, C., & Loomis, R. (1977). A new species of *Urosaurus* (Reptilia, Lacertilia, Iguanidae) from Baja California, Mexico. Journal of Herpetology, 11, 25–29.

Richardson, C. (1915). Reptiles of northwestern Nevada and adjacent territory. Proc. US Natl. Mus., 48, 403–435.

Riddle, B. R., Hafner, D. J., Alexander, L. F., & Jaeger, J. R. (2000). Cryptic vicariance in the historical assembly of a Baja California Peninsular Desert biota. Proceedings of the National Academy of Sciences, 97(26), 14438–14443. 10.1073/pnas.250413397

Riginos, C. (2005). Cryptic Vicariance in Gulf of California Fishes Parallels Vicariant Patterns Found in Baja California Mammals and Reptiles. Evolution, 59(12), 2678–2690. 10.1111/j.0014-3820.2005.tb00979.x

Schulte, J. A., Macey, J. R., & Papenfuss, T. J. (2006). A genetic perspective on the geographic association of taxa among arid North American lizards of the *Sceloporus magister* complex (Squamata: Iguanidae: Phrynosomatinae). Molecular Phylogenetics and Evolution, 39(3), 873–880. 10.1016/j.ympev.2005.04.033

Stamatakis, A. (2014). RAxML version 8: A tool for phylogenetic analysis and post-analysis of large phylogenies. Bioinformatics, 30(9), 1312–1313. 10.1093/bioinformatics/btu033

Stejneger, L. (1893). Annotated list of the reptiles and batrachians collected by the Death Valley Expedition in 1891, with descriptions of new species. North American Fauna, 7, 159– 228.

Stejneger, L. (1894). Description of *Uta mearnsi*, a new lizard from California. Proceedings of the United States National Museum, 17, 589–591.

Stock, J. M., & Hodges, K. V. (1989). Pre-Pliocene Extension around the Gulf of California and the transfer of Baja California to the Pacific Plate. Tectonics, 8(1), 99–115. 10.1029/TC008i001p00099

Sumarli, A., Hollingsworth, B. D., Valdez–Villavicencio, J. H., & Reeder, T. W. (2024). Phylogenomic data reveal cryptic diversity and deep phylogeographical structure within the common chuckwalla, *Sauromalus ater* (Squamata: Iguanidae). Biological Journal of the Linnean Society, 141(4), 572–588. 10.1093/biolinnean/blad104

Upton, D. E., & Murphy, R. W. (1997). Phylogeny of the side-blotched lizards (Phrynosomatidae: *Uta*) based on mtDNA sequences: support for midpeninsular seaway in Baja California. Molecular Phylogenetics and Evolution, 8(1), 104–113. 10.1006/mpev.1996.0392

Valdivia-Carrillo, T., García-De León, F. J., Blázquez, Ma. C., Gutiérrez-Flores, C., & González Zamorano, P. (2017). Phylogeography and Ecological Niche Modeling of the Desert Iguana (*Dipsosaurus dorsalis*, Baird & Girard 1852) in the Baja California Peninsula. Journal of Heredity, 108(6), 640–649. 10.1093/jhered/esx064

Van Denburgh, J. (1894). Descriptions of three new lizards from California and lower California, with a note on *Phrynosoma blainvillii*. Proceedings of the California Academy of Sciences, 4, 296–301.

Van Denburgh, J. (1895). Review of the herpetology of lower California. Proceedings of the California Academy of Sciences, 2, 77–163.

Van Denburgh, J. (1922). The reptiles of western North America, Vol 1: Lizards. Proceedings of the California Academy of Sciences, 1, 1–611.

Wiens, J. J., Hutter, C. R., Mulcahy, D. G., Noonan, B. P., Townsend, T. M., Sites, J. W., & Reeder, T. W. (2012). Resolving the phylogeny of lizards and snakes (Squamata) with extensive sampling of genes and species. Biology Letters, 8(6), 1043–1046. 10.1098/rsbl.2012.0703

Yarrow, H. C. (1882). Descriptions of new species of reptiles and amphibians in the United States National Museum. 10.5479/si.00963801.299.438

Zink, R. M. (2002). Methods in Comparative Phylogeography, and Their Application to Studying Evolution in the North American Aridlands. Integrative and Comparative Biology, 42(5), 953–959. 10.1093/icb/42.5.953

