## Supplementary figures and images for "Comparative phylogeography of lizards (Squamata: Phrynosomatidae) in Baja California and expansion of *Callisaurus draconoides* within the North American deserts"

### Supplemental Figure 1

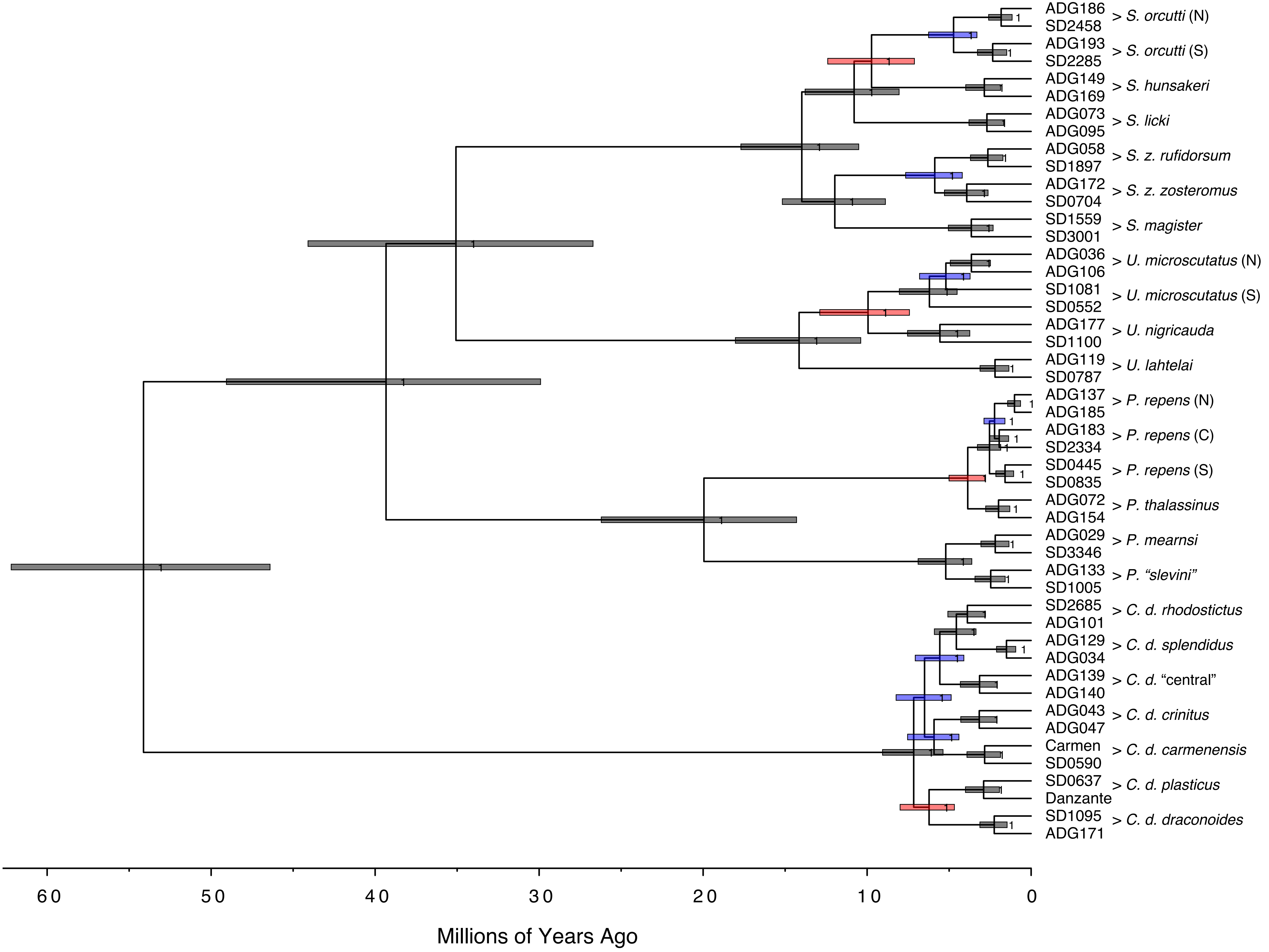

### Supplemental Figure 2

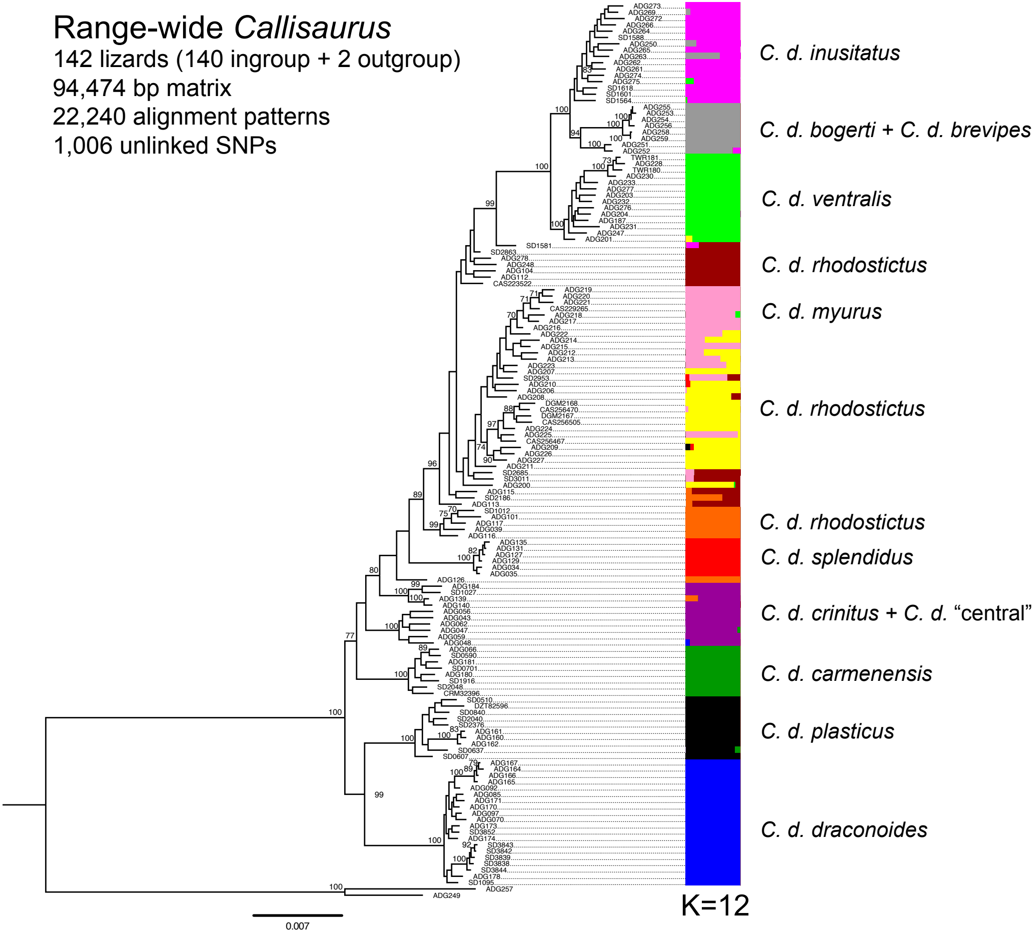
